# A vast evolutionarily transient translatome contributes to phenotype and fitness

**DOI:** 10.1101/2021.07.17.452746

**Authors:** Aaron Wacholder, Saurin Bipin Parikh, Nelson Castilho Coelho, Omer Acar, Carly Houghton, Lin Chou, Anne-Ruxandra Carvunis

## Abstract

Translation is the process by which ribosomes synthesize proteins. Ribosome profiling recently revealed that many short sequences previously thought to be noncoding are pervasively translated. To identify protein-coding genes in this noncanonical translatome, we combine an integrative framework for extremely sensitive ribosome profiling analysis, iRibo, with high-powered selection inferences tailored for short sequences. We construct a reference translatome for *Saccharomyces cerevisiae* comprising 5,400 canonical and almost 19,000 noncanonical translated elements. Only 14 noncanonical elements were evolving under detectable purifying selection. Surprisingly, a representative subset of translated elements lacking signatures of selection demonstrated involvement in processes including DNA repair, stress response and post-transcriptional regulation. Our results suggest that most translated elements are not conserved protein-coding genes and contribute to genotype-phenotype relationships through fast-evolving molecular mechanisms.

## Introduction

The central role played by protein-coding genes in biological processes has made their identification and characterization an essential project for understanding organismal biology. Over the past decade, the scope of this project has expanded as ribosome profiling (ribo-seq) studies have revealed pervasive translation of eukaryotic genomes.^1–4^ These experiments demonstrate that genomes encode not only the “canonical translatome”, consisting of the open reading frames (ORFs) identified as protein-coding genes in genome databases like RefSeq^5^, but also a large “noncanonical translatome” consisting of ORFs that are not annotated as genes. Despite lack of annotation, large-scale studies find that many noncanonical ORFs are translated to express stable proteins and show evidence of association with cellular phenotypes.^6–10^ Additionally, a handful of previously unannotated coding sequences, identified by RNA-seq or ribo-seq experiments, have now been characterized in depth, revealing that they play key roles in biological pathways and are important to organism fitness.^11–15^ Yet, these well-studied examples represent only a small fraction of the noncanonical translatome. Most noncanonical translation could simply be biologically insignificant “translational noise” resulting from the imperfect specificity of translation processes.^16–19^ Alternatively, thousands of missing protein-coding genes that contribute to phenotype and fitness could be hidden in the noncanonical translatome.

A common and powerful approach to identifying biologically significant genomic sequences is to look for evidence of selection.^20–22^ Many canonical genes were annotated on the basis of such evidence^23, 24^, and this approach has also been applied to noncanonical ORFs detected by ribo-seq.^25–28^ However, in the case of noncanonical translation, evolutionary analysis is often limited by a lack of sufficient statistical power to confidently detect selection. Most noncanonical ORFs are much shorter than canonical genes^7, 12, 29^, thus having fewer genetic variants that can be analyzed for evolutionary inference. As a result, short coding sequences are sometimes missed by genome-wide evolutionary analyses despite long-term evolutionary conservation.^13, 30^ It is especially challenging to detect selection among noncanonical ORFs that are evolutionarily novel, as a short evolutionary history also provides less time for enough genetic variants to accumulate the signatures that allow for statistically distinguishing selective from neutral evolution.^31^ Several young genes of recent *de novo* origin (i.e., coding genes that evolved from previously nongenic sequences) have been discovered from within the noncanonical translatome.^3, 32, 33^

In addition to the challenges short ORF length poses for detection of selection, it also poses challenges for unequivocal detection of translation in the first place. Microproteins are often missed by most proteomics techniques, though specialized methods have had some success.^9, 10, 34–36^ In ribo-seq data, the most robust evidence of translation comes from a pattern of triplet periodicity in reads corresponding to the progression of the ribosome across codons.^6, 37, 38^ Ribo-seq analysis methods are less capable of detecting translation of short ORFs, as they contain fewer positions to use to establish periodicity.^39^ The low expression levels of some noncanonical ORFs further increases the difficulties in identification.^3, 27^ Perhaps as a result of these limitations, less than half of the noncanonical ORFs detected as translated in humans are reproducible across studies.^31^

Here, we designed an approach to increase sensitivity in detection of both translation and selection among noncanonical ORFs. We address the challenges in detecting translation through the development of a ribo-seq analysis framework (iRibo) that identifies signatures of translation with high sensitivity and high specificity by integrating data across hundreds of experiments from many published studies. This facilitates detection of sequences that are short or poorly expressed. We address the challenges in detecting selection through a comparative genomics framework that analyzes translated sequences collectively across evolutionary scales within- and between-species.

We applied our approach to define a “reference translatome” for the model organism *Saccharomyces cerevisiae* and to characterize the biological significance of noncanonical ORFs. Using iRibo, we identified ∼19,000 noncanonical ORFs translated at high confidence and established the dependence of noncanonical translation on both genomic context and environmental condition. Using genomic data both within strains of *S. cerevisiae* and across budding yeast species^40, 41^, we identified a handful of undiscovered conserved genes within the yeast noncanonical translatome. However, we find that most of the yeast noncanonical translatome is evolutionarily young and of *de novo* origin, having emerged recently from noncoding sequence. These young ORFs differ greatly from conserved genes in their length, amino acid composition, and expression level, and show no signs of purifying selection. Nevertheless, we report experimental evidence based on fluorescent protein tagging and conditional loss-of-function fitness measurements showing that translation of evolutionarily young noncanonical ORFs can generate stable protein products and affect cellular phenotypes. We thus propose that much of the noncanonical translatome is composed of neither translational noise nor conserved genes, but rather of a distinct class of evolutionarily short-lived coding sequences with important biological implications. This “transient translatome” is larger than, and categorically distinct from, the conserved translatome made mostly of canonical protein-coding genes that have been studied for decades.

## Results

### An integrative approach to defining the translatome

We designed iRibo to detect translation events with high sensitivity and high specificity. High sensitivity is achieved through integration of ribo-seq data across hundreds of diverse experiments, which provides sufficient read depth for detection of translated ORFs that are short or weakly expressed. High specificity is achieved through the use of three nucleotide periodicity as the sole basis for translation inference. Three nucleotide periodicity corresponds to the progression of the ribosome codon by codon across a transcript, a dynamic unique to translation. Three nucleotide periodicity is therefore robust against false inference of translation from other sources of ribo-seq reads.^37, 38, 42^ High specificity is further achieved by controlling confidence levels using an empirical false discovery rate approach that relies on minimal modeling assumptions. iRibo consists of four components (**Figure 1A**). First, a set of “candidate” ORFs that could potentially be translated are identified in the genome. Second, reads from multiple ribo-seq experiments are pooled and mapped to these ORFs. Third, the translation status of each candidate ORF is assessed based on whether the reads mapping to the ORF exhibit a pattern of triplet nucleotide periodicity according to a binomial test. Finally, a list of translated ORFs is constructed with a specified false discovery rate, derived from applying the same translation calling method on a negative control set constructed to exhibit no genuine signatures of translation.

**Figure 1:**
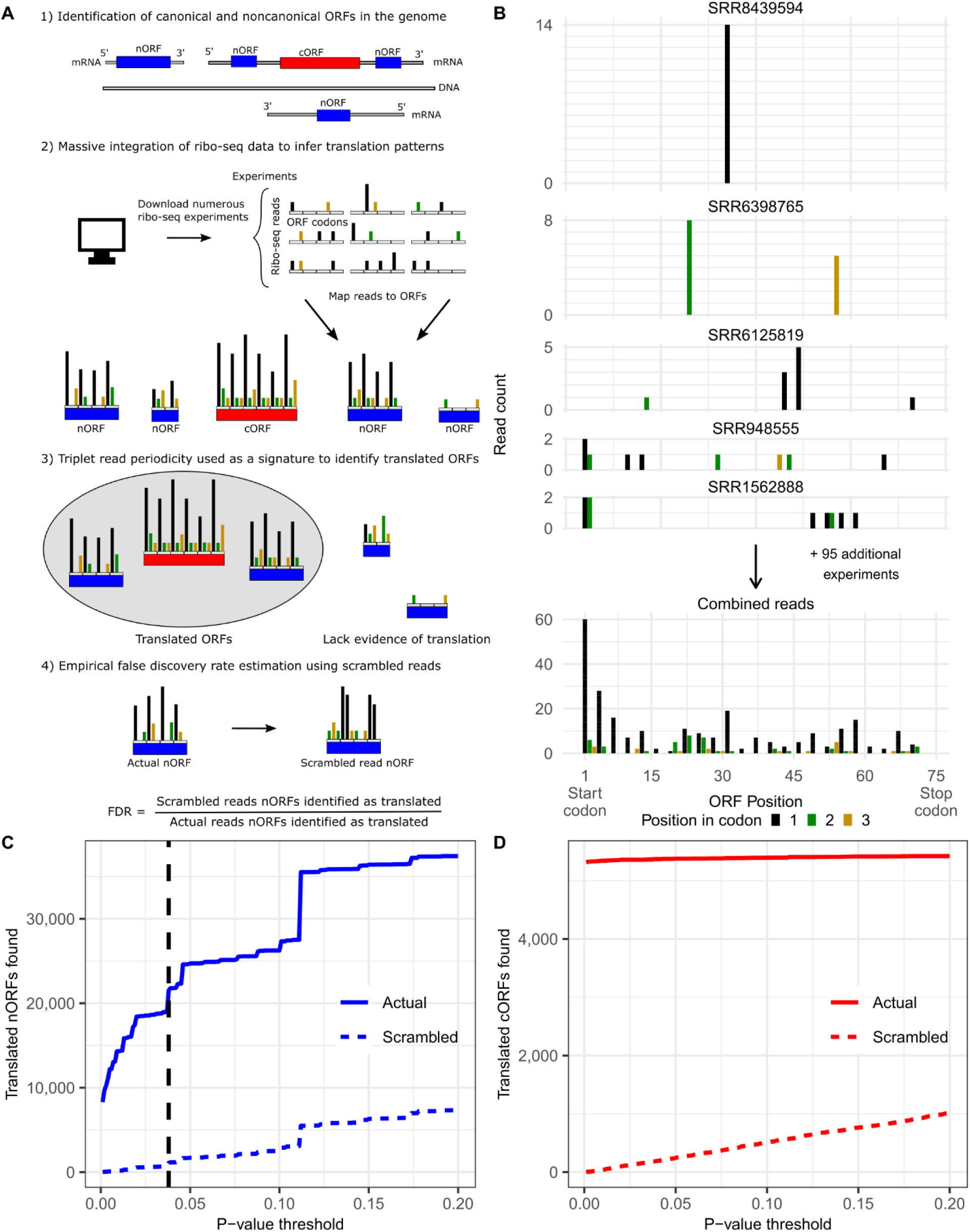
The iRibo framework enables detection of thousands of noncanonical translated sequences. **A)** The iRibo framework. 1) Candidate ORFs, both canonical (cORFs; red) and noncanonical (nORFs; blue), are identified in the genome. 2) Reads aggregated from published datasets are then mapped to these ORFs. 3) Translation is inferred from triplet periodicity of reads. 4) The false discovery rate is estimated by scrambling the ribo-seq reads of each ORF and then assessing periodicity in this scrambled set. **B)** iRibo identifies translated ORFs that are undetectable in any single experiment. Mapped ribo-seq reads (y-axis) across an example nORF located on chromosome II, 604674-604748 (x-axis). The top five graphs correspond to five individual experiments with reads mapping to the ORF while the bottom graph includes all reads integrated across all experiments. Reads are colored according to their position on the codon. **C)** iRibo identifies 18,953 translated nORFs at 5% false discovery rate. The number of nORFs found to be translated using iRibo (y-axis) at a range of p-value thresholds (x-axis) is shown as a solid blue line. Translation calls for a negative control set, constructed by scrambling the actual ribo-seq reads for each nORF, is also plotted (dashed blue line). The dashed vertical line indicates false discovery rate of 5% among nORFs. **D)** iRibo identifies 5,364 cORFs. The number of cORFs found to be translated using iRibo at a range of p-value thresholds, contrasted with negative controls constructed by scrambling the ribo-seq reads of each cORF.

iRibo can be applied to a set of ribo-seq experiments conducted under a single environmental condition to identify ORFs that are translated under that condition. Alternatively, iRibo can be deployed on a broader set of ribo-seq experiments conducted in many different contexts to construct a “reference translatome” consisting of all elements within a genome with sufficient evidence of translation.

We used iRibo to identify translated ORFs across the *S. cerevisiae* genome (**Supplementary Figure 1**). First, we constructed the set of candidate ORFs by collecting all genomic sequences at least three codons in length that start with ATG and end with a stop codon in the same frame. For ORFs overlapping in the same frame, only the longest ORF was kept. Each candidate ORF was classified either as canonical (cORF), if it was annotated as “verified,” “uncharacterized,” or “transposable element” in the Saccharomyces Genome Database (SGD)^43^ or as noncanonical (nORF), if it was annotated as “dubious,” “pseudogene,” or was unannotated. We excluded nORFs that overlap cORFs on the same strand. This process generated a list of 179,441 candidate ORFs: 173,868 nORFs and 5,573 cORFs. We assessed translation status for candidate ORFs using data from 412 ribo-seq experiments across 42 studies (**Supplementary Table 1, Supplementary Table 2**).

As expected, integrating data from many experiments allowed for identification of translated ORFs that would otherwise have too few reads in any individual experiment (**Figure 1B**). Setting a confidence threshold to ensure a 5% false discovery rate (FDR) among nORFs, we identified 18,953 nORFs (**Figure 1C**) as translated along with 5,364 cORFs (**Figure 1D**), for a total of 24,317 ORFs making up the yeast reference translatome. This corresponds to an identification rate of 99% for “verified” cORFs, 77% for “uncharacterized” cORFs, 37% for “dubious” nORFs and only 11% for unannotated nORFs (**Figure 2A**). Despite the low rate of identified translation, unannotated nORFs make up a large majority of translated es **(Figure 28).** In general, translated cORFs are much longer ( Figure2C) and translated at much ates **(Figure 2D)** than translated nORFs.

**Figure 2:**
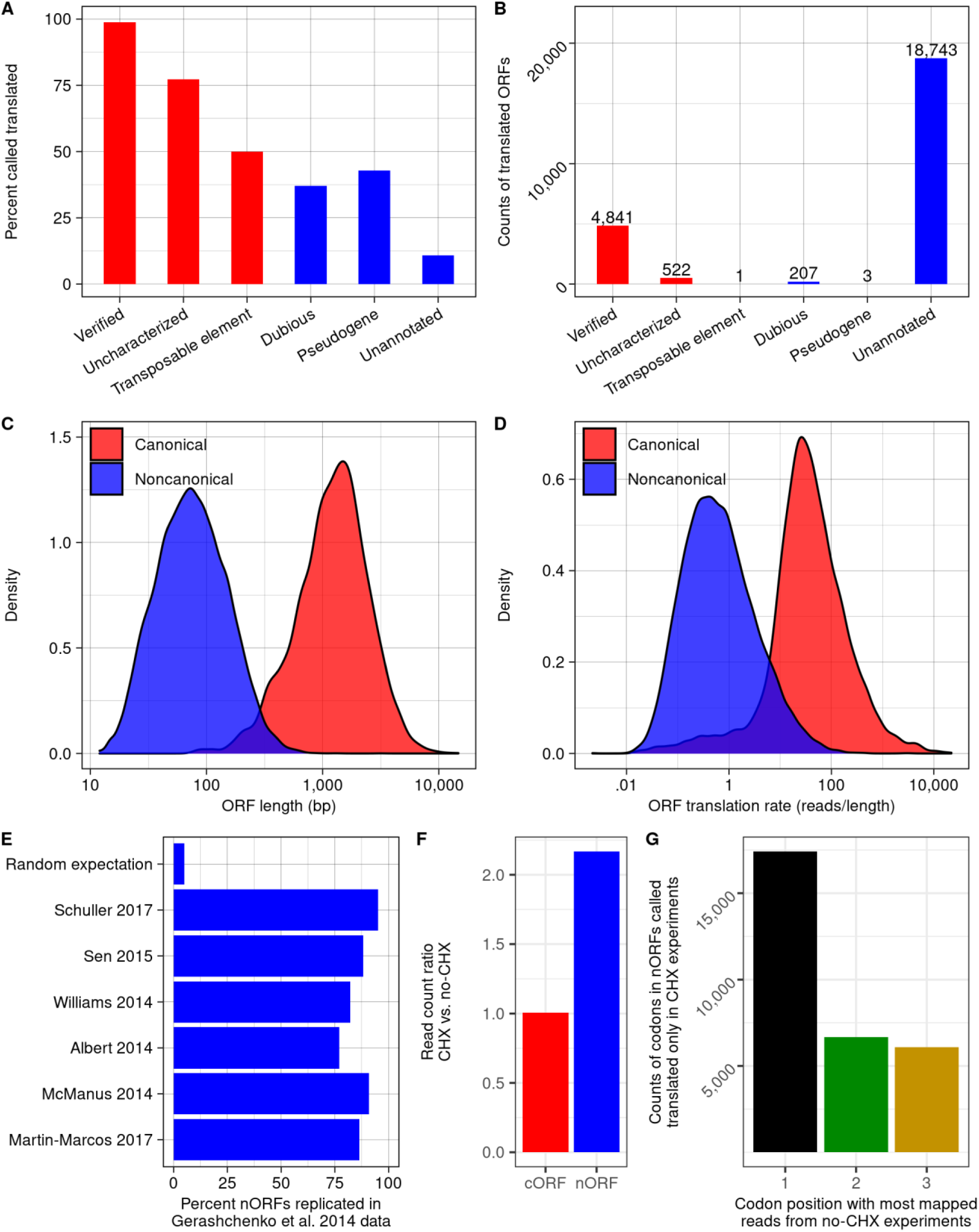
The noncanonical yeast translatome is larger than the canonical. **A)** A majority of cORFs, and a minority of nORFs, are translated. The percent of ORFs (y-axis) in each Saccharomyces Genome Database annotation class that are detected as translated by iRibo, with canonical classes indicated in red and noncanonical in blue. **B)** Unannotated nORFs make up a large majority of translated sequences. The number of ORFs (y-axis) of each annotation class (x-axis) that are detected using iRibo. **C)** nORFs are shorter than cORFs. ORF length distributions for translated cORFs and nORFs. **D)** nORFs are translated at lower rates than cORFs. Distribution of translation rate (in-frame ribo-seq reads per base) for translated cORFs and nORFs. **E)** Translation calls are highly reproducible. For six large studies (y-axis), the proportion of nORFs identified using reads from that study that are replicated using reads from the largest study, Gerashchenko et al. 2014^44^ (x-axis). Random expectation is the proportion that would be expected to replicate by chance. **F)** CHX facilitates detection of translated nORFs. Ratio of total ribo-seq read counts mapping to cORFs or nORFs in experiments with vs. without CHX treatment. Note that the same number of total reads are sampled from each condition. **G)** nORFs identified as translated only with CHX nevertheless show preference for the first codon position in its absence. Among nORFs identified as translated by iRibo only in the CHX condition, all codons among these nORFs are classed based on which of the three positions in the codon have the most reads from experiments without CHX.

To assess replicability in translation calls for nORFs, we applied iRibo separately to each of the largest individual studies by read count. We then counted, among the nORFs that could be inferred to be translated using only the reads in each study, how many were also found in the largest study, Gerashchenko and Gladyshev, 2014.^44^ For all studies, at least 75% of detected ORFs were also detected in the largest study (**Figure 2E**). In general, translation rates among ORFs were highly correlated among independent studies (**Supplementary Figure 2**). These observations demonstrate that noncanonical translation patterns are highly reproducible, suggesting that they are driven by regulated biological processes rather than technical artifacts or stochastic ribosome errors.

A large fraction of ribo-seq experiments use the translation elongation inhibitor cycloheximide (CHX). This drug is known to influence ribo-seq results in several ways.^44–46^ We therefore wished to specifically examine whether the size of the noncanonical translatome we identified could have been artificially inflated by CHX usage. To this aim, we compared translation signatures from experiments with (N=139) and without (N=170) CHX, randomly sampling the same number of reads from both groups of experiments. We observed a large enrichment in ribo-seq read counts among nORFs with CHX treatment (p < 10^-10^, Fisher’s exact test, **Figure 2F**), resulting in 56% more nORFs identified as translated (p < 10^-10^, Fisher’s exact test). The nORFs identified as translated only with CHX treatment nevertheless displayed a strong collective signal of triplet periodicity (i.e., preferential mapping to the first position in the codon) in experiments without CHX treatment when reads were aggregated across all such nORFs (**Figure 2G**). These results indicate that CHX treatment aids detection of translation events that also occur but are more difficult to detect without CHX.

### Noncanonical translation patterns depend on genomic and environmental context

We examined to what extent translation of nORFs depends on genomic context. We classified nORFs as: upstream nORFs (uORFs) located on the 5’ untranslated regions of transcripts containing cORFs; downstream nORFs (dORFs) located on the 3’ untranslated regions of transcripts containing cORFs; intergenic nORFs that do not share transcripts with cORFs (independent); nORFs antisense to a cORF and located entirely within the bounds of that cORF (antisense full overlap); nORFs overlapping the boundaries of a cORF on the opposite strand (antisense partial overlap) (**Figure 3A**). Additionally, for nORFs sharing a transcript with an RNA gene, the nORF was classified based on the type of RNA gene. The transcripts used for these classifications were derived from the TIF-seq data collected by Pelechano et al. 2014^47^, which provide transcript start and end sites.

**Figure 3:**
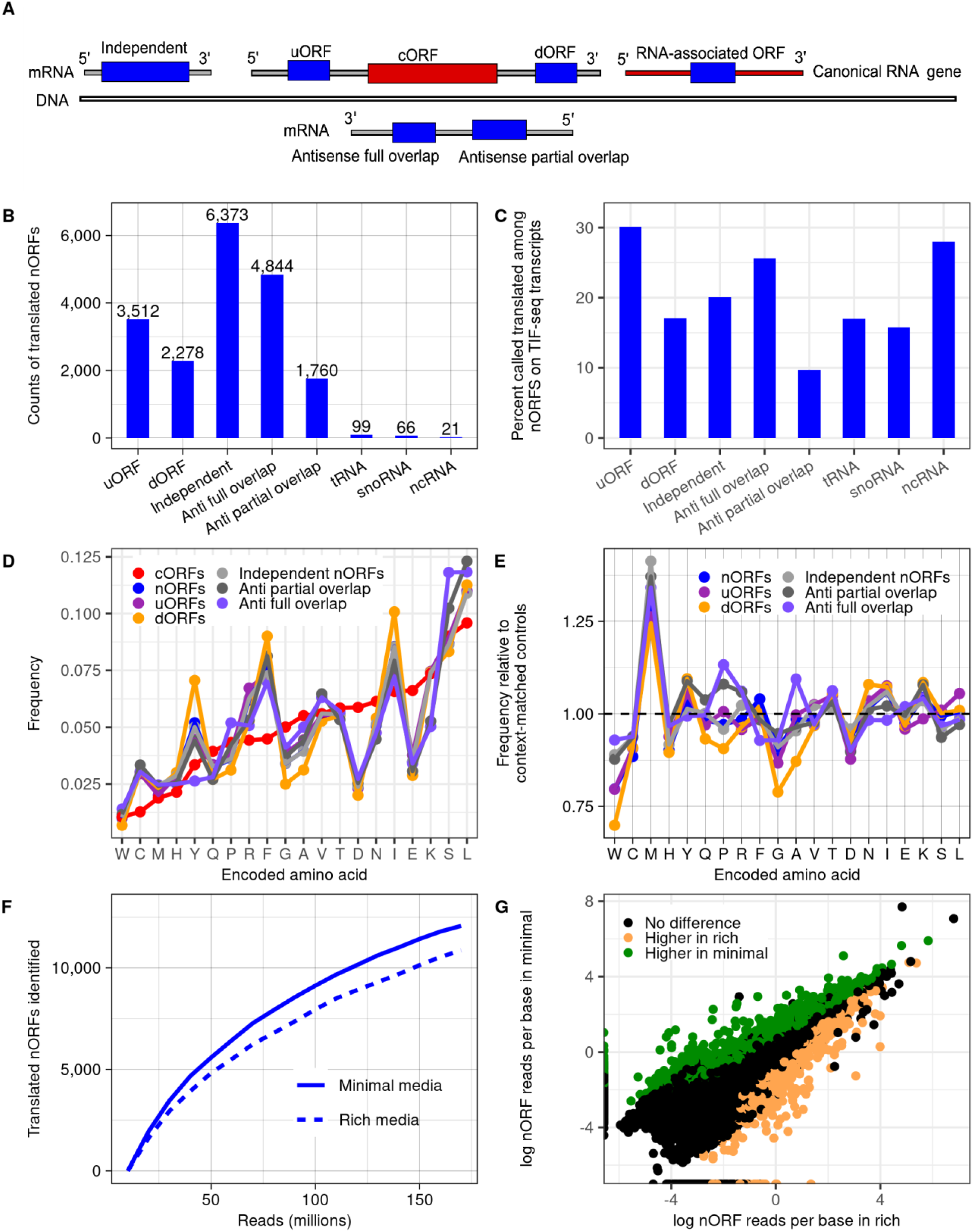
Noncanonical translation patterns depend on both genomic and environmental context. **A)** Potential genomic contexts for nORFs in relation to nearby canonical genes. Transcripts are defined from published TIF-seq data47. **B)** Counts of translated nORFs identified by iRibo (y-axis) in each considered genomic context (x-axis), determined by which elements share a transcript with the nORF and its position within the transcript. For nORFs that share a transcript with RNA genes, the annotation of the RNA gene is specified. **C)** Proportion of nORFs detected as translated by iRibo (y-axis) in each genomic context considered, among nORFs completely covered by a TIF-seq transcript (x-axis). **D)** Amino acid composition of translated nORFs differs from that of translated cORFs and depends on genomic context. Amino acid frequencies among predicted protein products of translated nORFs in each genomic context and of cORFs. The start codon methionine is excluded from frequency estimates. **E)** Amino acid composition of translated nORFs is similar to that of context-matched controls. For each genomic context, the amino acid frequency of translated nORFs relative to that of length-matched untranslated nORFs in that same context. The start codon methionine is excluded from frequency estimates. **F)** More nORFs are identified as translated in minimal than rich media. Number of translated nORFs identified (y-axis) for experiments on yeast grown in either minimal (SD, solid line) or rich media (YPD, dashed line) at a range of read depths (x-axis). For each read depth, reads are sampled at random from experiments in each condition. **G)** For each nORF called translated by iRibo in minimal media (SD), rich media (YPD), or both, the log reads per base in each condition is indicated. Total read count in each condition was held constant by randomly sampling reads from YPD experiments until the read count in SD experiments was matched. nORFs with significantly more reads in one condition than the other are colored, green for SD and brown for YPD. Lists of nORFs with significantly different translation rates were obtained as follows: p-values for differential translation of each nORF were calculated from Fisher’s exact test on in-frame ribo-seq reads mapping to the ORF in each condition and a 5% FDR was set using the Benjamini-Hochberg approach.^48^ An nORF had to be detected as translated in a condition by iRibo to be identified as more highly translated in that condition.

Most nonoverlapping translated nORFs were independent (6,373, 52%) and around 47% shared a transcript with a cORF, including 3,512 uORFs and 2,278 dORFs, while 1.5% (186) shared a transcript with an annotated RNA gene (**Figure 3B**). Among antisense nORFs, 73% (4,844) overlapped fully with the opposite-strand gene while 27% (1,760) overlapped partially.

We next calculated the frequency at which candidate nORFs were identified as translated for each genomic context (**Figure 3C**); for purposes of comparison, we considered only those nORFs fully contained within a TIF-seq transcript. Consistent with prior research^49^, uORFs were translated at significantly higher rates than other classes, with 30% of considered uORFs found to be translated compared to only 17% of dORFs (p < 10^-10^, Fisher’s Exact Test) and 20% of independent nORFs (p < 10^-10^, Fisher’s Exact Test). nORFs antisense to cORFs and only partially overlapping them were translated at the lowest rate of any context, with a rate of 10% compared to 26% for fully overlapping antisense nORFs (p < 10^-10^, Fisher’s Exact Test).

The amino acid frequencies of the proteins expressed from translated nORFs differ greatly from those of cORFs and depend on genomic context (p<10^-10^ for any comparison between cORF amino acid frequencies and nORF frequencies in a given context, chi-square test; **Figure 3D**). The translation products of nORFs present a large excess of cysteine, phenylalanine, isoleucine, arginine, and tyrosine and deficiency in alanine, asparagine, glutamic acid, and glycine relative to cORFs. Notably, aside from arginine, the amino acids with large excess in nORFs relative to cORFs are all hydrophobic. Amino acid frequencies of nORFs appear to largely reflect underlying DNA sequence composition biases that differ between the distinct genomic contexts. Indeed, within each genomic context, amino acid frequencies of translated nORF are generally similar (with less than 15% difference in frequency) to that of length- and context-matched nORFs that lack evidence of translation, though they do show significant differences (p < 10^-10^ for all contexts, chi-square test; **Figure 3E**). The most striking differences include a large excess of methionine residues and a deficiency in tryptophan and glycine residues among translated nORFs compared to the untranslated control group.

In addition to genomic context, we assessed how environmental context affects noncanonical translation. To this aim, we leveraged the power of iRibo to construct separate datasets of nORFs found translated in rich media (YPD) or in nutrient-limited minimal media (SD) (**Supplementary Table 3**). Previous research has reported an increase in detected noncanonical translation events relative to canonical translation events in response to starvation.^1,3^ Consistent with these results, more nORFs were identified as translated in minimal than in rich media at equal read counts (**Figure 3F**). Furthermore,

2968 nORFs were supported by a significantly higher number of in-frame reads in minimal media than rich media while the converse was true for only 1265 nORFs (5% FDR, Fisher’s exact test with Benjamini-Hochberg procedure^48^; **Figure 3G**). These results suggest that starvation conditions may increase noncanonical translation, or alternatively that noncanonical translation is less affected by the general translation inhibition that occurs in starvation conditions.^50^ Either way, these results support the hypothesis that nORF translation is regulated in response to changing environments.

### Two translatomes, transient and conserved

Given the large numbers of nORFs translated in the yeast genome, we next sought to assess the biological significance of this translation by determining the extent to which these nORFs are evolving under selection. We assessed selection acting on nORFs, as well as on cORFs for purpose of comparison, across three evolutionary scales. At the population level, we analyzed 1011 distinct *S. cerevisiae* isolates sequenced by Peter et al. 2018.^40^ At the species level, we compared *S. cerevisiae* ORFs to their orthologs in the *Saccharomyces* genus, a taxon consisting of *S. cerevisiae* and its close relatives.^51^ To detect long term evolutionary conservation, we looked for homologs of *S. cerevisiae* ORFs among 332 budding yeast genomes (excluding *Saccharomyces*) in the subphylum *Saccharomycotina* collected by Shen et al. 2018.^41^

The power to detect selection on an ORF depends on the amount of genetic variation in the ORF available for evolutionary inference, which in turn depends on its length, the density of genetic variants across its length, and the number of genomes available for comparison. Given that many translated nORFs are very short (**Figure 2C**), we employed a two-stage strategy to increase power for detecting signatures of selection. First, we investigated selection in a set of “high information” ORFs for which we have sufficient statistical power to potentially detect selection. Second, we investigated the remaining “low information” ORFs in groups to quantify collective evidence of selection (**Figure 4A**). Group level analysis increases power to detect the presence of selection but does not enable identification of the individual ORFs under selection. The “high information” set consisted of the ORFs that 1) have homologous DNA sequence in at least four other *Saccharomyces* species and 2) have a median count of nucleotide differences between the *S. cerevisiae* ORF and its orthologs of at least 20. We found these criteria are sufficient to distinguish ORFs evolving under strong purifying selection (**Supplementary Figure 3**). Under this definition, 9,440 translated ORFs that do not overlap a different cORF (henceforth “nonoverlapping ORFs”, including 4,248 nORFs, and 5,192 cORFs) and 3,022 ORFs that overlap a cORF on the opposite strand (“antisense ORFs”, including 2,962 nORFs and 60 cORFs) were placed in the “high information” set.

**Figure 4:**
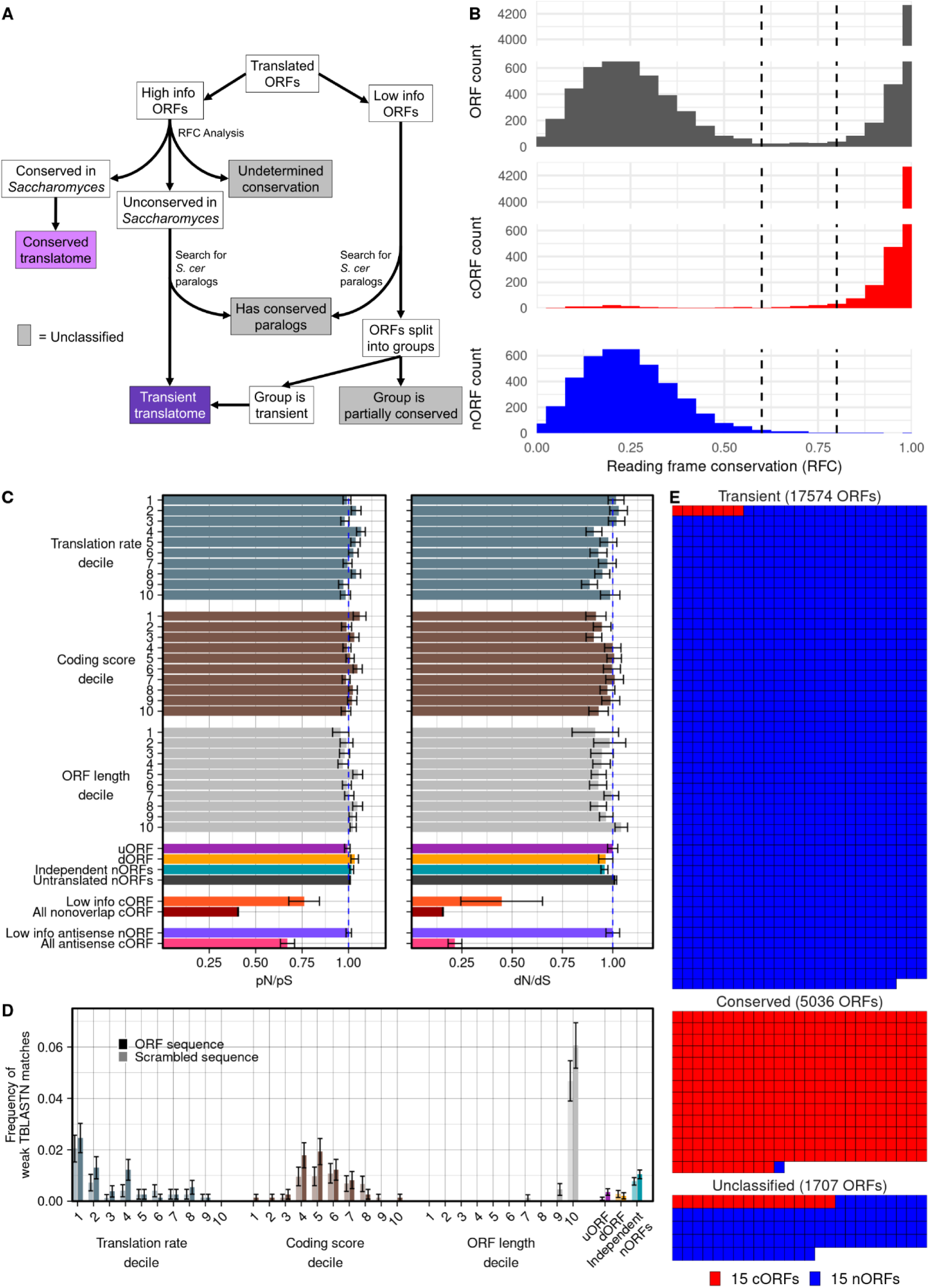
Two distinct translatomes: transient and conserved. **A)** Selection inference analyses conducted on low-information and high-information ORFs to classify them as evolutionarily conserved, transient, or unclassified. **B)** A bimodal distribution of reading frame conservation (RFC) among high information translated ORFs. The distribution of RFC (x-axis), indicating how well reading frame of the ORF is conserved in the Saccharomyces genus, is shown for all translated high information ORFs (top), only cORFs (middle) and only nORFs (bottom). See Methods for details. Dashed lines separate RFC < 0.6 and RFC > 0.8, the thresholds used to distinguish ORFs preserved or not preserved by selection. **C)** No evidence of purifying selection acting on low information nORFs. pN/pS and dN/dS ratios are shown for each group of ORFs. Low information nonoverlapping nORFs that lack a conserved homolog are divided into deciles of translation rate (in-frame ribo-seq reads per base), coding score, or ORF length, and into three genomic contexts. Untranslated nORFs are the set of all nORFs in the genome not called as translated by iRibo. Low information nonoverlapping cORFs are assembled into a single group, with the set of all nonoverlapping cORFs shown for comparison. Low information antisense nORFs were also assembled into a single group, with the set of all antisense cORFs shown for comparison. pN/pS is calculated from variation at each ORF codon among *S. cerevisiae* isolates.^40^ dN/dS is calculated among all codons that share the same frame between *S. cerevisiae* ORFs and aligned orthologous ORFs in *S. paradoxus*. Note that the displayed pN/pS and dN/dS values are not averages of these ratios among ORFs. Rather, synonymous and nonsynonymous variants among all ORFs in each class are counted, and a single ratio is calculated from the summed counts. Error bars indicate standard errors estimated from bootstrapping. The dashed blue line indicates a ratio of one, the expected ratio under neutral evolution. **D)** No evidence of distant homology for low information nORFs. The frequency of nORFs with weak TBLASTN matches (10^-4^ < e-value < .05) in each group of nORFs (dark bars) and negative controls (light bars) consisting of the sequences of the nORFs of each group randomly scrambled. Error bars indicate standard errors estimated from bootstrapping. **E)** ORFs that are translated yet evolutionarily transient make up 72% of the yeast reference translatome. The components of the translatome (transient, conserved, unclassified) are represented with area proportional to frequency. Each box represents sets of 15 ORFs.

We attempted to detect purifying selection in the high information set within the *Saccharomyces* genus and within the *Saccharomycotina* subphylum. For the *Saccharomyces* analysis, we adapted reading frame conservation (RFC), a sensitive approach developed by Kellis et al. 2003^20^ to distinguish ORFs evolving under selection from other ORFs in the yeast genome. RFC is an index ranging from 0 to 1 that indicates how well reading frame is conserved between an ORF in a given species (here, *S. cerevisiae*) and its orthologous sequences in related species (other species in the *Saccharomyces* genus). An RFC value of 1 indicates perfect agreement of reading frame, such that all bases that make up the first nucleotide in a codon in the *S. cerevisiae* ORF also make up the first nucleotide in a codon in each orthologous ORF. An RFC value of 0 indicates that all bases in the *S. cerevisiae* ORF align to bases with a different within-codon position in orthologous ORFs, or that the aligned bases exist outside of any ORF. We found a bimodal distribution of RFC among nonoverlapping ORFs in the yeast translatome, considering cORFs and nORFs together: 53.3% have RFC above 0.8 and 45.5% have RFC less than 0.6, with only 1.2% of ORFs intermediate between these values (**Figure 4B)**. The bimodal distribution of RFC among translated ORFs is similar to the bimodal distribution observed among all candidate ORFs, regardless of translation status (**Supplementary Figure 4A**), as observed previously by Kellis et al. 2003.^20^

The modes of the distribution largely correspond to annotation status, with 96.7% of cORFs having RFC > 0.8 and 98.5% of nORFs having RFC < 0.6. This pattern holds when evaluated only in the last 100 bp of ORFs, suggesting that it is not affected by potential incorrect inference of nORF start positions (**Supplementary Figure 4B**). The clean separation between well-conserved and poorly-conserved ORFs indicate that most high-information ORFs can be straightforwardly classified into one of the two groups, and thus nearly all high-information nonoverlapping nORFs can be placed in the poorly-conserved class. High RFC among antisense ORFs does not demonstrate selection on the ORF itself, as it might be caused by selective constraints on the opposite-strand gene, but low RFC still indicates lack of purifying selection. A majority of antisense translated nORFs (64.1%) have RFC <0.6, indicating that most are not preserved by selection across the genus (**Supplementary Figure 4C**). Overall, we find no evidence for purifying selection acting on nORFs on a large scale.

In light of the general correspondence between annotation and conservation, the exceptions are of interest: 110 cORFs had RFC < 0.6, and 13 nonoverlapping unannotated nORFs had RFC > 0.8. To further assess conservation among these two sets of ORFs, we performed a BLAST analysis (using both BLASTP and TBLASTN with default parameters) to search for homologs of each ORF among the budding yeast genomes assembled by Shen et al. 2018.^41^ Among the 110 cORFs with low RFC, 101 also had no detected homology to other *S. cerevisiae* genes or any budding yeast genome outside of *Saccharomyces*, indicating that these are likely annotated ORFs of recent *de novo* origin. For the 13 nORFs with high RFC, several additional lines of evidence suggest that these are indeed evolving under purifying selection (**Table 1)**. For nine of the thirteen, we identified a homolog among budding yeast genomes outside of the *Saccharomyces* genus by either a BLASTP or TBLASTN search. The existence of a homolog in a distantly related species indicates that the ORF existed in the common ancestor of *S. cerevisiae* and that distant species, implying long-term preservation of the ORF by purifying selection in both lineages. We also performed pN/pS analysis for each ORF on *S. cerevisiae* isolates and dN/dS analysis for each ORF among the *Saccharomyces* genus species (**Table 1**). A pN/pS or dN/dS ratio significantly below 1 indicates purifying selection on the ORF amino acid sequence among *S. cerevisiae* strains or among *Saccharomyces* genus species, respectively, while a ratio above 1 indicates positive selection. By these measures, two ORFs showed significant evidence of purifying selection by pN/pS and three by dN/dS (**Table 1**). Thus, a small number of nORFs appear to be evolving under selection, indicating significant biological roles.

**Table 1:** Properties of well-conserved nORFs. Systematic name refers to either the systematic name annotated on SGD, or the name assigned here according to SGD conventions. BLASTP and TBLASTN e-values are the minimum BLASTP or TBLASTN e-value observed in a search of the ORF against the yeast genomes assembled by Shen et al.^41^, excluding those in the *Saccharomyces* genus. BLAST coverage is the length of the segment that aligns to the best identified homolog (lowest e-value) in the BLAST search. RFC is reading frame conservation of the ORF among species in the Saccharomyces genus. Length is the length of the ORF in nucleotides. The pN/pS ratio is obtained from nucleotide variation in the ORF among the 1011 S. cerevisiae strains assembled by Peter et al.^40^; significant values below 1 indicate purifying selection. The dN/dS ratio was obtained from nucleotide variation in the ORF among Saccharomyces genus species; significant values below 1 indicate purifying selection. Translation percentile indicates the percentage of nORFs with a lower ribo-seq read count per codon than the indicated ORF.

| Systematic Name | Coordinates | BLASTP e-value | TBLASTN e-value | RFC | BLAST coverage (nt) | Length (nt) | pN/pS (p-value) | dN/dS (p-value) | Translation percentile |
| --- | --- | --- | --- | --- | --- | --- | --- | --- | --- |
| YBL029W-B<sup>a</sup> | chrII:164192-164368 | 6.5 x 10<sup>-4</sup> | 8.0 x 10<sup>-3</sup> | 0.82 | 107 | 177 | 1.65 (.33) | 0.88 (.68) | 67 |
| YBL014W-A<sup>a</sup> | chrII:196737-196889 | 4.1 x 10<sup>-5</sup> | 1.0 x 10<sup>-4</sup> | 1 | 116 | 153 | 0.47 (.11) | 0.14 (3.46 x 10<sup>-12</sup>) | 86 |
| YBR085W-B<sup>a</sup> | chrII:417494-417556 | 1 | 1 | 0.86 | 0 | 63 | 0.72 (.48) | 1.26 (.62) | 58 |
| YBR268W-A<sup>a</sup> | chrII:741844-742005 | 1 | 1 | 0.99 | 0 | 162 | 0.61 (.15) | 0.35 (3.18 x 10<sup>-7</sup>) | 97 |
| YBR292W-A<sup>a</sup> | chrII:786745-786903 | 1.9 x 10<sup>-7</sup> | 5.0 x 10<sup>-3</sup> | 0.96 | 146 | 159 | 0.72 (.43) | 0.57 (.0026) | 83 |
| YER186W-A<sup>a</sup> | chrV:565603-565800 | 6.0 x 10<sup>-6</sup> | 1 | 0.92 | 143 | 198 | 0.55 (.02) | 1.0 (1) | 97 |
| YGL262W-A<sup>a</sup> | chrVII:4663-4872 | 1 | 1.0 x 10<sup>-3</sup> | 0.88 | 113 | 210 | 0.96 (.86) | 1.0 (1) | 86 |
| YGR238W-A<sup>a</sup> | chrVII:969015-969089 | 1 | 1 | 0.87 | 0 | 75 | 0.20 (.01) | 1.18 (.74) | 94 |
| YBL049C-A<sup>a</sup> | chrII:126330-126461 | 8.3 x 10<sup>-5</sup> | 6.0 x 10<sup>-4</sup> | 0.84 | 92 | 132 | 1.36 (.79) | 1.5 (.22) | 75 |
| YBL026C-A<sup>a</sup> | chrII:169634-169870 | 6.8 x 10<sup>-12</sup> | 9.0 x 10<sup>-10</sup> | 0.88 | 116 | 237 | 1.30 (.6) | 0.87 (.42) | 99.96 |
| YJR107C-A<sup>a</sup> | chrX:628457-628693 | 3.8 x 10<sup>-8</sup> | 3.0 x 10<sup>-18</sup> | 0.99 | 161 | 237 | 0.39 (.005) | 1.42 (.13) | 99.91 |
| YLR349C-A<sup>a</sup> | chrXII:828276-828338 | 1 | 1 | 0.81 | 0 | 63 | 0.30 (.02) | 0.73 (.24) | 73 |
| YNR062C-A<sup>a</sup> | chrXIV:745640-745792 | 5.0 x 10<sup>-14</sup> | 5.0 x 10<sup>-13</sup> | 0.89 | 110 | 153 | 0.65 (.44) | 1.49 (.15) | 44 |
| YBR012C | chrII:259147-259566 | 6.51 x 10<sup>-59</sup> | 1x10<sup>-16</sup> | 0.70 | 120 | 420 | .62 (.1) | .50 (.039) | 92 |

<sup>a</sup>We assigned this unannotated ORF a systematic name based on SGD conventions.

We next assessed selection among the full set of nORFs (both high and low information) at the subphylum scale, searching for addition nORFs that exhibited long term conservation and thus purifying selection. Towards this end, we searched for distant homologs of all translated nonoverlapping *S. cerevisiae* nORFs using TBLASTN against budding yeast genomes in the *Saccharomycotina* subphylum, excluding species in the *Saccharomyces* genus. After excluding matches that appeared non-genic or pseudo-genic (**Supplementary Figure 5**) we identified a single additional nORF with both distant TBLASTN matches and recent signatures of purifying selection (dN/dS = 0.5, p=.039 for test of difference from 1.0): YBR012C, annotated as “dubious” on SGD. Thus, combining the 13 nORFs that appeared conserved by RFC analysis and the single additional nORF found using TBLASTN, we identified 14 translated nORFs that show evidence of preservation by purifying selection (**Table 1**).

To analyze collective evidence of selection among “low information” ORFs, we first divided low information nonoverlapping nORFs (7,855 nORFs, after excluding those with homology to conserved *S. cerevisiae* cORFs) according to properties that we expected to be potentially associated with selection: rate of translation (as measured by ribo-seq reads mapped to the first position within codons divided by the length of the ORF), coding score^28, 52^ (a measure of sequence similarity to annotated coding sequences), ORF length, and genomic context. For each group, we calculated the pN/pS ratio among 1,011 *S. cerevisiae* isolates^40^ and the dN/dS ratio based on alignments of the *S. cerevisiae* ORFs with their orthologous DNA sequence in *S. paradoxus*. We also analyzed low information nonoverlapping cORFs (22 cORFs) in the same manner. For low information antisense nORFs (3642 nORFs; only 2 cORFS fell in this category and were not analyzed), we calculated the pN/pS and dN/dS ratios restricted to substitutions that were synonymous on the opposite-strand cORF.^53, 54^ Unlike the RFC, dN/dS and pN/pS analyses conducted above on individual high information ORFs, these analyses were conducted by aggregating substitutions among all low information ORFs in each group to assess evidence for selection (i.e., a ratio significantly different from 1) within the group as a whole. We expected that, if low information nORFs were evolving under selection, then more highly translated ORFs, longer ORFs, and ORFs with coding scores more similar to conserved genes, would be enriched in biologically relevant nORFs and thus show stronger signatures of selection. Low information nonoverlapping cORFs did show collective pN/pS and dN/dS ratios significantly below 1, indicating that some ORFs in this group are evolving under purifying selection (**Supplementary Table 4**, **Figure 4C**). In contrast, for all groups of low information nORFs examined, we observed no significant difference in the pN/pS or dN/dS ratio from 1, providing no evidence for either purifying or positive selection (**Supplementary Table 4, Figure 4C**).

Finally, we assessed collective evidence of long-term evolutionary conservation in each group. To do this, we calculated the frequency of weak TBLASTN matches (e-values between 10^-4^ and .05, above our threshold for homology detection at the individual level) of ORFs in each group to the *Saccharomycotina* subphylum genomes outside of *Saccharomyces* as compared to a negative control set consisting of scrambled sequences of the ORFs in each group. Applying this strategy to the full set of 362 nonoverlapping cORFs that lacked TBLASTN matches outside *Saccharomyces* at the e-value < 10^-4^ level, we found a large excess of weak matches relative to controls (p=.0001, Fisher’s exact test; **Supplementary Figure 6**), demonstrating the ability of this approach to detect faint signals of homology within a group of ORFs. However, we identified no significant difference in the frequency of weak TBLASTN hits between any nonoverlapping nORF group and scrambled controls (**Figure 4D**), nor among nonoverlapping nORFs overall (p>.05, Fisher’s exact test). The lack of a significant result does not exclude the possibility that a small subset of short conserved nORFs could be lost in the noise of a much larger set of nORFs without distant homology. However, our TBLASTN, dN/dS and pN/pS analyses altogether indicate that ORFs evolving under strong purifying selection are not a major component of the yeast noncanonical translatome.

Overall, our analyses distinguish two distinct yeast translatomes: a conserved, mostly canonical translatome with intact ORFs preserved by selection; and a mostly noncanonical translatome where ORFs are not preserved over evolutionary time. This distinction is rooted in evolutionary evidence rather than annotation history. We thus propose to group the translated ORFs that showed neither evidence of selection nor homology to conserved ORFs in our high-information and low-information sets as the “transient translatome.” The “transient translatome” designation indicates membership in a set of ORFs that are expected to exist in the genome for only a short time on an evolutionary scale, though we cannot be certain that any particular translated ORF that currently exists in the yeast genome will be rapidly lost. The transient translatome includes 4,051 nonoverlapping and 1,923 antisense nORFs identified as not preserved by selection using RFC analyses and having no conserved homologs, along with 86 nonoverlapping and 15 antisense cORFs (total 101) matching the same criteria. Also included are 7,855 nonoverlapping and 3,644 antisense nORFs that lack sufficient information to analyze at the individual level but were found to show no selective signal in group-level analyses. Together, this set of 17,574 ORFs that are translated yet likely evolutionarily transient makes up 72% of the yeast reference translatome (**Figure 4E**).

### Transient cORFs are representative of the transient translatome overall

By general theory and practice in evolutionary genomics, the lack of selective signal suggests that the transient translatome does not meaningfully contribute to fitness.^55^ Surprisingly, however, 101 cORFs belong to the transient set, suggesting that some transient ORFs have phenotypes. To assess whether these cORFs are representative of the transient translatome overall, we compared their evolutionary and sequence properties with those of transient “dubious” nORFs (annotated but presumed nonfunctional) and transient unannotated nORFs. We found transient cORFs, transient dubious nORFs and transient unannotated nORF to all have pN/pS ratios indistinguishable from 1.0 (**Figure 5A**), providing no evidence for purifying selection. Similarly, the average nucleotide diversity (mean number of nucleotide differences per site between pairs of isolates) of transient cORFs was indistinguishable from that of transient nORFs or untranslated controls, and much higher than that of conserved cORFs (**Figure 5B**). In addition, no class of transient ORFs showed differences from each other in RFC between *S. cerevisiae* and *S. paradoxus* (**Figure 5C**), rate of translation (**Figure 5D**) or coding score (**Figure 5E)**.

**Figure 5:**
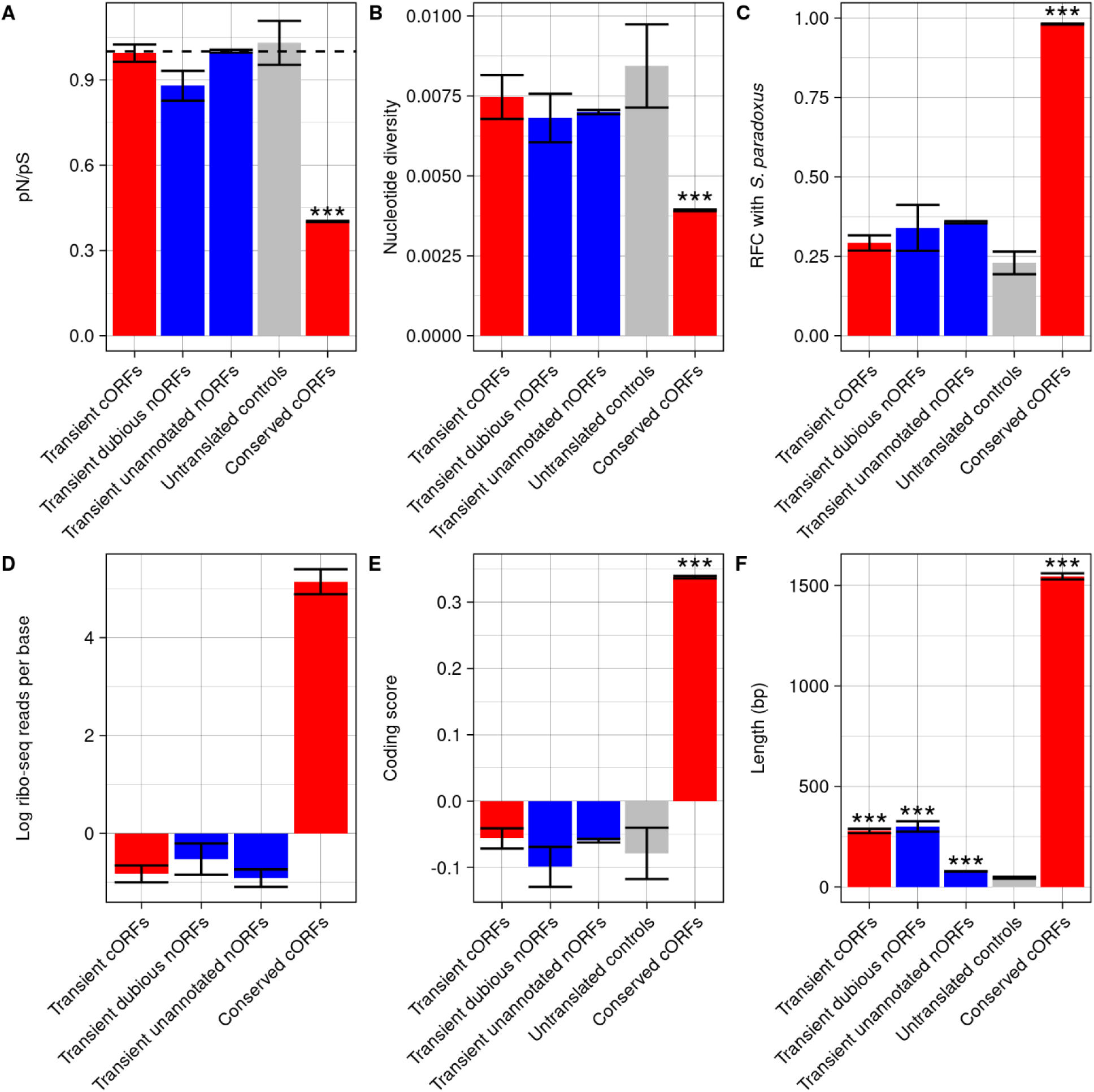
Canonical and noncanonical transient ORFs are similar except for their length. Properties of transient cORFs (n=86), transient dubious nORFs (n=25), transient unannotated nORFs (n=12,160), untranslated controls (n=100) and conserved cORFs (n=5130). Untranslated controls consist of nonoverlapping ORFs that would be grouped in the transient class (RFC <.6) but are not inferred to be translated based on ribo-seq evidence. Conserved cORFs are nonoverlapping cORFs with RFC >.8. All groups are restricted to nonoverlapping ORFs. Error bars represent standard error. Stars indicate significant differences from untranslated controls by permutation test: P-value <.001: ***. **A)** pN/pS values for each group among *S. cerevisiae* strains. **B)** Average nucleotide diversity (π) among each group. **C)** Average reading frame conservation between *S. cerevisiae* and *S. paradoxus* ORFs. **D)** Average ribo-seq reads per base (logged), considering only in-frame reads. Unannotated nORFs and untranslated controls are sampled to match the length distribution of transient cORFs. **E)** Coding scores for each group. **F)** ORF lengths in nucleotides for each group.

The only distinguishing property between classes of transient ORFs was their length: annotated transient cORFs and transient “dubious” nORFs are much longer on average than unannotated transient nORFs (**Figure 5F**). This is a consequence of the history of annotation of the *S. cerevisiae* genome, where a length threshold of 300 nt was set for annotation of ORFs.^56, 57^ The sharp 300 nt threshold is still clearly reflected in annotations. For example, genome annotations include 96% of nonoverlapping transient ORFs in the 300-400 nt range (55/57), but only 4% in the 252-297 nt range (4/101). Given that transient nORFs resemble transient cORFs in all respects besides length, we hypothesized that numerous never-studied transient nORFs are just as likely to have phenotypes as transient cORFs.

### Transient ORFs are detected in the cell and mediate diverse phenotypes

To gain further insights into the potential biological roles of transient ORFs, we examined published reports about annotated ORFs (transient cORFs and transient dubious nORFs) in the *S. cerevisiae* experimental literature and performed additional experiments to investigate transient unannotated nORFs. We examined whether transient ORF products could be detected experimentally, whether they affect phenotypes, and whether they interact with specific biological pathways.

We first assessed whether the proteins encoded by transient ORFs can be detected in the cell. We examined the CYCLoPs database^58, 59^, the C-SWAT tagging library^60^, and the YeastRGB database^61^, which contain collections of fluorescently tagged proteins expressed from their native promoters and terminators, including both cORFs and dubious nORFs. Together these studies detected expression of a fluorescent protein product for 90 of 93 (97%) transient cORFs tested, along with 37 of 41 (90%) transient dubious nORFs tested (**Figure 6A**). For comparison, we C-terminally tagged 21 highly expressed unannotated transient nORFs with mNeonGreen at their endogenous locus and examined their expression using microscopy. We detected 8 of 21 tagged nORF proteins (38%) (**Figure 6A-B, Supplementary Figure 7**). Thus, translation of tagged proteins can be detected for both annotated and unannotated transient ORFs.

**Figure 6:**
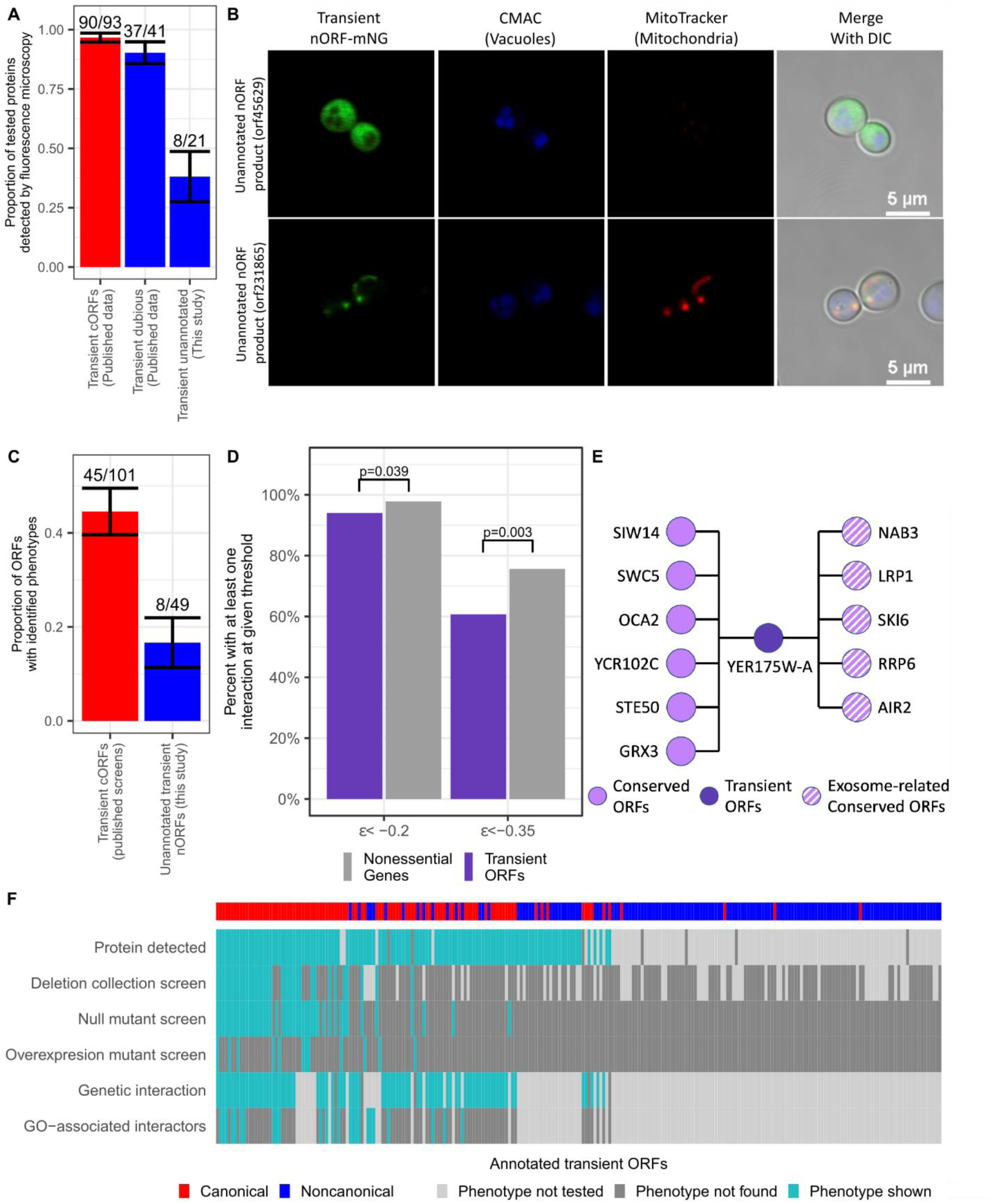
Transient nORFs and cORFs can be detected in the cell and exhibit phenotypes. **A)** Transient in this study. Error bars indicate standard error of the proportion. **B)** Tagged unannotated transient nORFs show varied sub-cellular localizations. Microscopy images of unannotated transient nORFs taken at 100X. Left panel shows the expression of the nORFs tagged with mNeonGreen, middle panels the dyes CMAC Blue and MitoTracker Red for mitochondria and vacuoles identification, respectively, and the right panel the merge all the above channels with DIC. Top panel show the nORF (orf45629) with a cytosolic expression and the bottom panel the nORF (orf231865) with expression localizing to the mitochondria. **C)** Loss of transient nORFs can affect phenotype despite lack of evolutionary conservation The proportion of deletion mutants with reported loss-of-function phenotypes in two groups: transient cORFs in published deletion mutant screens, and transient nORFs assayed in this study. Reported phenotypes in published data was taken from literature associated with each ORF on SGD. In this study, deleterious deletion mutant phenotypes were identified from a high-throughput colony fitness screen in six stress conditions using a 5% FDR threshold. **D)** Transient ORFs engage in epistatic relationships. The percent of transient ORFs and nonessential genes with at least one genetic interaction at given threshold are shown. Differences between groups were tested using Fisher’s exact test. **E)** Genetic interactions of the transient ORF YER175W-A. Five interactors are related to exosome (striped circles). **F)** Presence of phenotypes among annotated transient ORFs. “Protein detected” indicates that the ORF product was found in either the C-SWAT or CYCLoPs database. Phenotypes of deletion collection, deletion and overexpression screens were taken from reported findings in the yeast experimental literature (**Supplementary Table 5**). “Genetic interaction” indicates a statistically significant genetic interaction with Ɛ< -0.2, and “GO-associated interactors” indicates a GO enrichment was found among significant interactors at 5% FDR.

We next examined the evidence that transient ORFs affect phenotype. Five transient cORFs have been studied in depth. Two of these, *MDF1*^62^ and YBR196C-A^63^, have been previously described as having emerged *de novo* from non-genic sequences. *MDF1* inhibits the mating pathway in favor of vegetative growth^62, 64^ and YBR196C-A is an ER-located transmembrane protein whose expression is beneficial under nutrient limitations.^65^ The remaining three have been experimentally characterized, although their evolutionary properties were not analyzed in the corresponding studies: *HUR1* plays an important role in non-homologous end-joining DNA repair^66^; *YPR096C* regulates translation of *PGM2*^67^; *ICS3* is involved in copper homeostasis.^68^ These cases demonstrate that some transient ORFs do affect phenotypes and have the potential to play important biological roles.

To determine whether transient cORFs that are not well described also affect phenotypes, we examined all literature listed as associated with the ORF on SGD. Many of these transient cORFs have direct evidence of phenotype (**Supplementary Table 5**). Of 101 transient cORFs, 45 were reported to have deletion mutant phenotypes (i.e., a phenotype observed when the ORF is deleted) and 12 to have overexpression phenotypes. Overall, we found phenotypes reported in the literature for 50 of 101 transient cORFs (50%).

As unannotated transient nORFs have not been systematically investigated for phenotype, we sought to experimentally determine whether these ORFs too might have deletion mutant phenotypes. We thus conducted a deletion mutant screen of 49 unannotated transient nORFs selected for high translation rate and to avoid intersecting cORFs, annotated ncRNAs, or promoters (200 bp upstream of canonical genes). We fully deleted the nORF using homologous recombination and each strain was assayed for colony growth in seven conditions. Eight nORF deletion mutant strains showed deleterious phenotypes in at least one condition at a 5% FDR (**Figure 6C**, **Supplementary Table 6**). Thus, loss of transient nORFs, as with cORFs, can affect phenotype despite lack of evolutionary conservation.

To begin to understand the specific biological processes in which transient ORFs might be involved, we leveraged the large yeast genetic interaction network assembled in Costanzo et al. 2016.^69^ This dataset includes 75 non-overlapping transient cORFs and 9 non-overlapping dubious transient nORFs. Genetic interaction strength, Ɛ, measures the difference between the observed fitness of a strain in which two genes are deleted and the expected fitness given the fitness of the two single gene deletion strains; a negative value of high magnitude suggests that the two mutated genes are involved in related processes. Of the 84 transient ORFs in the dataset, 79 (94%) have at least one negative genetic interaction at the high-stringency cut-off defined by Costanzo et al.^69^ (Ɛ<-0.2 and p-value<0.05) and 51 (61%) have synthetic lethal interactions (Ɛ<-0.35 and p-value<0.05) as defined in that study (**Figure 6D**). This was only a slightly lower rate than for conserved non-essential ORFs, 98% of which had negative interactions at the high stringency cut-off and 76% of which had synthetic lethal interactions. At the high stringency threshold, 27 transient ORFs were found to interact with groups of related genes enriched in specific gene ontology (GO) terms (5% FDR; **Supplementary Table 7**). For example, the interactors of YER175W-A are associated with the GO category “cryptic unstable transcript (CUT) metabolic processes” with high confidence, and five of its eleven interactors are components or co-factors of the exosome (**Figure 6E**), indicating likely involvement in CUT degradation or a closely related post-transcriptional regulation pathway. Other enrichments included diverse processes such as “mating projection tip” or “Golgi sub-compartment”. In contrast, when we applied GO enrichment analysis to the full set of genes that interact with any transient ORF, no significant enrichment was observed. These results suggest that transient ORFs in general do not participate in one shared biological process, but rather are involved in a wide variety of cellular processes.

Overall, we uncovered evidence that 131 of 250 (53%) annotated transient ORFs have at least one indicator of biological significance (detection of a protein product, a reported phenotype in a screen, or a genetic interaction in the Costanzo et al. 2016^69^ network) (**Figure 6F**). Additionally, we demonstrate that unannotated transient ORFs encode proteins that can be detected in the cell (38% of tested in this study) and influence cellular fitness when deleted (17% of tested in this study). Given that this class has received almost no study compared to the great number of experiments that have been conducted on cORFs, the number of transient ORFs with biological relevance may be substantially larger than that which has been annotated.

A limitation on much of the experimental evidence available on deletion mutant phenotypes is that most deletion mutant and genetic interaction screens are based on a full gene replacement strategy in which the entire ORF is lost, leaving the possibility that some deletion phenotypes could be caused by loss of a ncRNA or a DNA regulatory element located at the same position as the ORF rather than loss of the ORF translation (**Figure 7A**). To examine this possibility, we constructed a set of strains where the ORF start codon ATG was replaced with an AAG codon while keeping the rest of the ORF intact. This set included three transient cORFs that have previously been characterized on the basis of overexpression or full deletion mutants, *ICS3*^68^, YPR096C^67^, and YBR196C-A^65^, along with four transient nORFs that showed strong deleterious phenotypes in our full ORF deletion screen (**Supplementary Table 8**; *HUR1* and *MDF1* were not tested because they overlap other cORFs). Each deletion strain was tested in seven environmental conditions. The single nucleotide ATG→AAG mutation caused significantly reduced colony size for all three transient cORFs tested and for three of four transient nORFs tested in at least one condition (**Figure 7B**). We gave these three nORFs systematic names YDL204W-A, YGR016C-A, and YNL040C-A. The remaining nORF, YDR073C-A, showed a weak beneficial phenotype from the ATG→AAG mutation in some conditions, as did two other nORFs, YGR016C-A and YNL040C-A. The largest growth reductions were observed from disabling translation in YDL204W-A: this strain reached only 64% of wildtype growth in hydroxyurea and 63% in high salt concentration, with a smaller reduction to 94% growth in rich media (YPDA). These growth defects were also observed in a liquid growth setting (**Figure 7C-E**). To confirm that these phenotypes were caused by loss of the YDL204W-A protein rather than *cis* effects at the locus, we expressed the intact YDL204W-A ORF from a plasmid in the ATG→AAG mutant strain. Plasmid expression of the ORF fully restored the wildtype phenotype in the mutant strains (**Figure 7F-H**), providing further evidence that blocking YDL204W-A translation causes a loss of function phenotype mediated by loss of the encoded protein.

**Figure 7:**
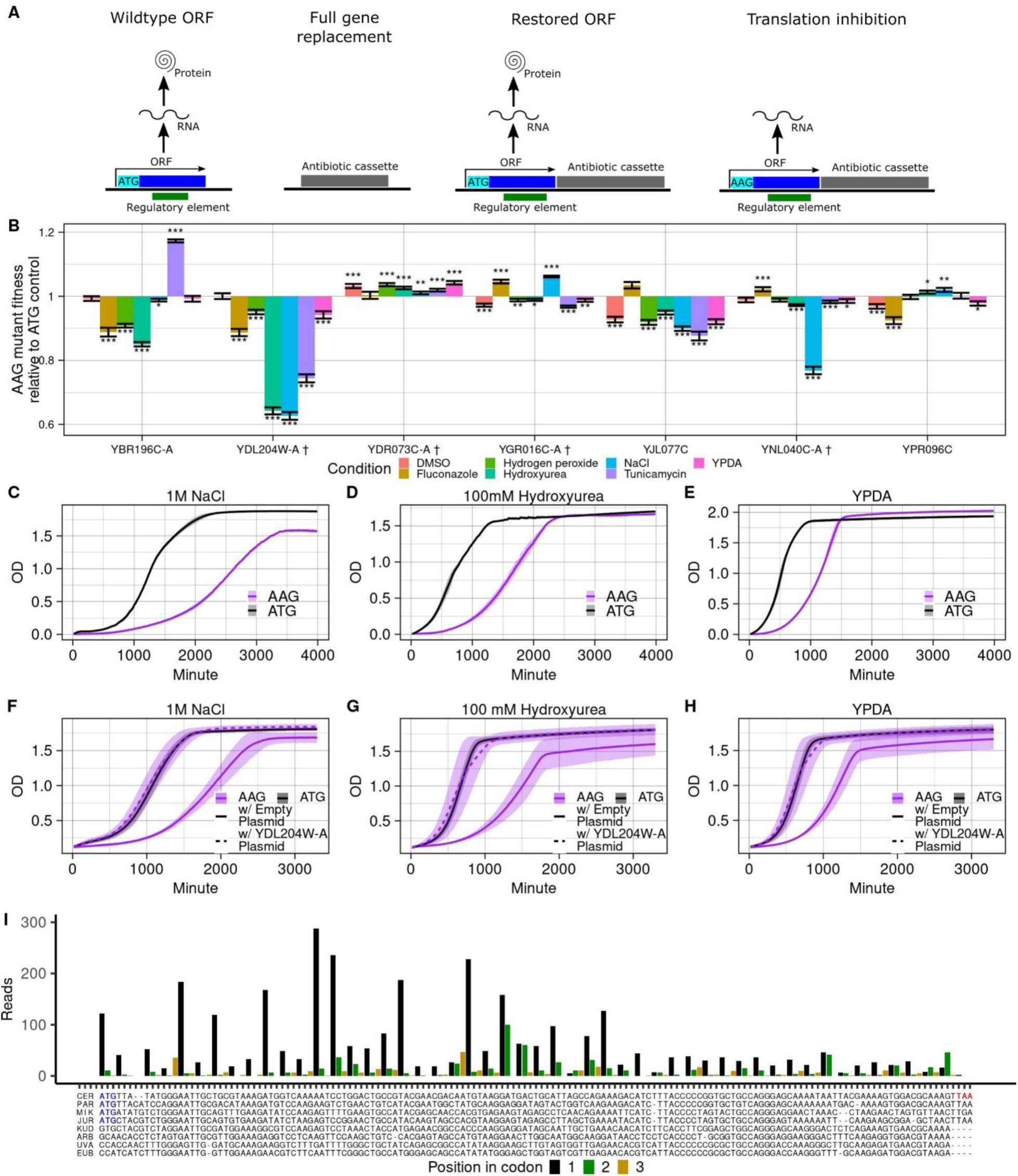
Translation inhibition of transient ORFs causes strong phenotypes. **A)** A two-step strategy for inhibiting nORF translation. An ORF may overlap a DNA regulatory element or an RNA with a noncoding function (Wildtype ORF), both of which are disrupted in a gene replacement strategy in addition to the loss of translation (Full gene replacement). This creates ambiguity in interpreting comparisons between deletion mutants and wildtype strains. Following a deletion screen using gene replacement, we used a second round of homologous recombination to restore either the full ORF (Restored ORF) or an ORF with its start codon mutated from ATG to AAG (Translation inhibition). As these mutants differ only by this single base, the specific effects of translation inhibition can be inferred. **B)** Inhibiting translation of transient ORFs triggers colony growth phenotypes. The fitness of AAG mutants (translation inhibition) is shown for seven transient ORFs under stress conditions (colors). Fitness is assessed by comparing colony size between AAG mutants and ATG controls (restored ORFs). A cross symbol after the ORF names indicates unannotated nORFs assigned systematic names in this study. Relative fitness values significantly different from one are indicated as follows: *p<.05 **p<.01 ***p<.001. **C-E)** Deleterious impact of inhibiting translation of transient nORF YDL204W-A in a liquid growth assay. Liquid growth curve of a strain in which YDL204W-A translation is inhibited by mutating its start codon (AAG) and a strain with the initial codon as ATG in: 1M NaCl (C), 100mM hydroxyurea (D), and YPDA (E), with three technical replicates for each strain. **F-H**) Expression from plasmid restores wildtype growth to YDL204W-A start codon mutants. Liquid growth curves of an attempted rescue of the YDL204W-A AAG mutant by expressing intact YDL204W-A from a plasmid. The AAG start codon mutants were transformed with either an empty plasmid or a plasmid expressing the intact ORF; the ATG controls were transformed with an empty plasmid. All strains were then assayed for growth in liquid media in either 1M NaCl (F), 100 mM hydroxyurea (G) or YPDA (H) with three technical replicates each. The shaded area covers 1 SD from the mean OD value among replicates. **I)** YDL204W-A is translated and not conserved. Top: ribosome profiling reads mapped by iRibo to YDL204W-A show triplet periodicity. Bottom: alignment of the YDL204W-A ORF against homologous DNA in the *Saccharomyces* genus.

In our translation dataset, YDL204W-A has a translation rate at the top percentile among transient ORFs (**Figure 7I**), higher than 10% of cORFs. Comparing its sequence to the homologous region of other *Saccharomyces* genus species, only *S. paradoxus* and *S. mikatae* have a homologous start codon, but a 2bp insertion in *S. cerevisiae* results in a frameshift such that little of the ORF is shared in any other species (**Figure 7I**); thus, this ORF has a reading frame conservation score of only 0.2 (**Table 2**). The other transient ORFs with phenotypes induced by an ATG→AAG mutation also showed no signs of selection **(Table 2)**. Thus, our results exemplify the potential for unannotated coding sequences with no evident evolutionary conservation to affect cellular phenotypes and fitness.

**Table 2:** Evolutionary properties of transient ORFs with phenotypes induced by inhibiting translation. The pN/pS ratio is obtained from nucleotide variation in the ORF among the 1011 S. cerevisiae strains assembled by Peter et al.^40^ TBLASTN was run for each ORF against genomes in the subphylum *Saccharomycotina*, excluding the genus *Saccharomyces*, with an e-value threshold of 10^-4^.

| ORF Name | Reading frame conservation | pN/pS (p-value) | TBLASTN matches |
| --- | --- | --- | --- |
| YBR196C-A | .29 | 1.34 (0.65) | 0 |
| YDL204W-A<sup>a</sup> | .20 | 1.25 (0.83) | 0 |
| YGR016C-A<sup>a</sup> | .29 | 0.66 (0.36) | 0 |
| YJL077C | .21 | 0.74 (0.19) | 0 |
| YNL040C-A<sup>a</sup> | .38 | 0.97 (1.00) | 0 |
| YPR096C | .20 | 1.39 (0.47) | 0 |

<sup>a</sup>We assigned this unannotated ORF a systematic name based on SGD conventions.

## Discussion

Since the advent of ribosome profiling, it has been evident that large parts of eukaryotic genomes are translated outside of canonical protein-coding genes^1^, but the nature and full significance of this translation has remained elusive. To facilitate study of this noncanonical translatome, we developed iRibo, a framework for integrating ribosome profiling data to sensitively detect ORF translation across a variety of environmental conditions. The iRibo framework can be applied to any species and set of candidate ORFs of interest. Here, we deployed iRibo to map a high confidence yeast reference translatome almost five times larger than the canonical translatome. This resource can serve as the basis for further investigations into the yeast noncanonical translatome, including the prioritization of nORFs for experimental study.

We designed iRibo to be highly sensitive at detecting patterns of triplet periodicity through the genome, but there are some limitations to our strategy. We focused exclusively on ORFs with AUG start codons and therefore missed the non-AUG codons that are sometimes used as starts.^70^ Similarly, we did not consider ORFs overlapping canonical genes in a different frame on the same strand, though some such nORFs are known to be translated.^71, 72^ Finally, candidate ORFs were selected as the longest ORF in any reading frame, which means the true boundaries of identified ORFs could be shorter than described. We expect these limitations to cause underestimation of the number of translated nORFs, suggesting that the true count is even larger than identified here.

We used the iRibo yeast reference translatome to address a fundamental question: to what extent does the noncanonical translatome consist of conserved coding sequences that were missed in prior annotation attempts? In a thorough evolutionary investigation, we identified 14 translated nORFs that show evidence of being conserved under purifying selection. Only one of these ORFs, YJR107C-A, appears to have been previously described^34^, though it was not annotated on Saccharomyces Genome Database at the time of our analysis. Thus, even a genome as well-studied as *S. cerevisiae*’s contains undiscovered conserved genes, likely missed in prior analyses due to difficulties in analyzing ORFs of short length. These 14 nORFs are, however, the exception: the great majority of translated nORF show no signatures of selection, comprising a large pool of evolutionarily transient translated sequences.

The yeast genome thus encodes two translatomes, one conserved, one transient. The conserved translatome consists of coding sequences that are preserved by strong purifying selection and usually have a long evolutionary history. They tend to be relatively long, well expressed, and with sequence properties highly distinct from noncoding sequences. The transient translatome, by contrast, is evolutionarily young, of recent *de novo* origin from previously noncoding sequence and still similar to noncoding sequences in nucleotide composition. Evolving in the absence of strong purifying selection, transient translated ORFs appear to be frequently lost to disrupting mutations, only to be replaced by other transient translated ORFs upon translation-enabling mutations. Despite these profound differences, transient translated ORFs, like conserved ones, can affect the phenotype and fitness of the organism. Several well-characterized coding sequences unique to *S. cerevisiae*, such as *HUR1*^66^ and *MDF1*^62^, play key roles in biological processes through encoding lineage-specific proteins that physically interact with conserved proteins. Additionally, around 100 transient ORFs are annotated as coding genes and have therefore been extensively screened; a majority express stable proteins and many have known loss-of-function phenotypes. Their genetic interaction patterns suggest involvement in a wide array of specialized cellular processes. Our experiments revealed that disabling the start codons of unannotated transient translated ORFs can cause large fitness reductions in stress conditions. The strength of the fitness reduction observed was highly dependent on the stressor applied in the environment, suggesting again specialized cellular roles. In some cases, disabling the start codon resulted in growth increases, perhaps indicating that disabling translation saved the cell energy.

Our work adds to the growing research on the roles noncanonical coding play across many species, including humans.^7, 73^ We note that “noncanonical” is not a coherent biological category, as it simply indicates the class of sequences that have not been annotated in genome databases. We demonstrate that the division between “canonical” and noncanonical” translation in *S. cerevisiae* corresponds largely, but not perfectly, to a biological division between transient and conserved. It is this biological division that is fundamental: the 101 yeast canonical ORFs classified as transient have sequence and evolutionary properties nearly identical to noncanonical transient ORFs, except for sequence length, and should be placed in the same category. We can thus reclassify the translatome according to biology rather than annotation history.

It is perhaps surprising that a coding sequence can affect organism phenotype despite showing no evidence of selection. However, this result is consistent with evidence from the field of *de novo* gene birth. Species-specific coding sequences have been characterized in numerous species.^32^ For example, Xie et al. 2019^74^ identified a mouse protein contributing to reproductive success that experienced no evident period of adaptive evolution. Sequences that contribute to phenotype without conservation have also been described outside of coding sequences. Regulatory sequences, such as transcription factor binding sites, are a mix of relatively well-conserved elements and elements that are not preserved even between close species^75^; it is plausible that translated sequences also show such a division. There are several explanations for why translated ORFs may lack detectable signatures of selection. Most transient ORFs are expressed at much lower levels than canonical genes, and therefore may have minimal effects on phenotype. For those that do have large and beneficial effects in some environmental conditions, these may be balanced by deleterious effects in other conditions. Moreover, selection may occur, and be biologically important, below the limits of detectability for the genomic approaches we used. Our findings do not imply an absence of selective forces in shaping the patterns of noncanonical translation. Rather, the particular selective environment favoring expression of these sequences may be too short-lived to detect selection using traditional comparative genomics approaches. Previous research, such as the proto-gene model of *de novo* gene birth^3^, have proposed that recently emerged translated ORFs serve as an intermediary between noncoding sequences and mature genes. Our results add to the evidence that these ORFs provide many potential phenotypes from which selection could preserve beneficial ones for the long term.^65^ Still, the observation that even ORFs with phenotypes lack evidence of conservation at the population level suggests that there are important filters that prevent the vast majority of recently emerged translated ORFs, even those with beneficial phenotypes, from evolving into mature genes that are preserved over long evolutionary time. The primary significance of the great majority of transient translated ORFs is in their biological activity over their short lifespans.

The yeast reference translatome resource we constructed with iRibo is meant to facilitate community efforts to decipher the specific physiological implications of transient translated ORFs. Our proof-of concept analyses of subcellular localization, genetic interactions and ATG->AAG mutants suggest involvement in diverse cellular processes and pathways. It is important to note that some transient translatome phenotypes may be mediated by a protein product, by the process of translation itself, or both. Translation of both uORFs^76^ and dORFs^77^ can affect expression of nearby genes. Translation also plays a major role in the regulation of RNA metabolism through the nonsense-mediated decay pathway.^78, 79^ Dissection of the molecular mechanisms mediating transient translatome phenotypes is an exciting area for future research.

Our results indicate that the yeast noncanonical translatome is neither a major reservoir of conserved genes missed by annotation, nor mere “translational noise.” Instead, many translated nORFs are evolutionarily novel and likely affect the biology, fitness, and phenotype of the organism through species-specific molecular mechanisms. As transient ORFs differ greatly in their evolutionary and sequence properties from conserved ORFs, they should be understood as representing a distinct class of coding element from most canonical genes. Nevertheless, as with conserved genes, understanding the biology of transient ORFs is necessary for understanding the relationship between genotype and phenotypes.

## Acknowledgments

We thank Dr. Emmanuel Doram Levy’s at the Weizmann Institute of Science for sharing the fluorescence intensity data displayed in YeastRGB. We thank Dr. Benjamin Dubreuil for the helpful discussion over YeastRGB data. We thank Dr. Allyson O’Donnell for her help in microscopy image acquisition. We thank Drs. Craig Kaplan and Nikolaos Vakirlis for helpful discussions of an earlier preprinted version of this manuscript. This work was supported by funds provided by the Searle Scholars Program to A.-R.C., the National Science Foundation grant MCB-2144349 to A.-R.C., and the National Institute of General Medical Sciences of the National Institutes of Health grants R00GM108865 and DP2GM137422

(awarded to A.-R.C.).

## Author contributions

Conceptualization, A.W. and A.-R.C. Methodology, A.W., A.-R.C., S.B.P., N.C.C., O.A. Investigation, A.W., N.C.C., S.B.P., O.A., C.H., L.C. Writing – Original Draft, A.W., S.B.P., O.A., N.C.C. Writing – Review & Editing, A.W., A.-R.C., S.B.P., N.C.C., O.A., C.H., L.C. Supervision, A.-R.C.

## Declaration of interests

A.-R.C. is a member of the scientific advisory board for Flagship Labs 69, Inc (ProFound Therapeutics).

## Supplementary figure legends

**Supplementary Figure 1:**
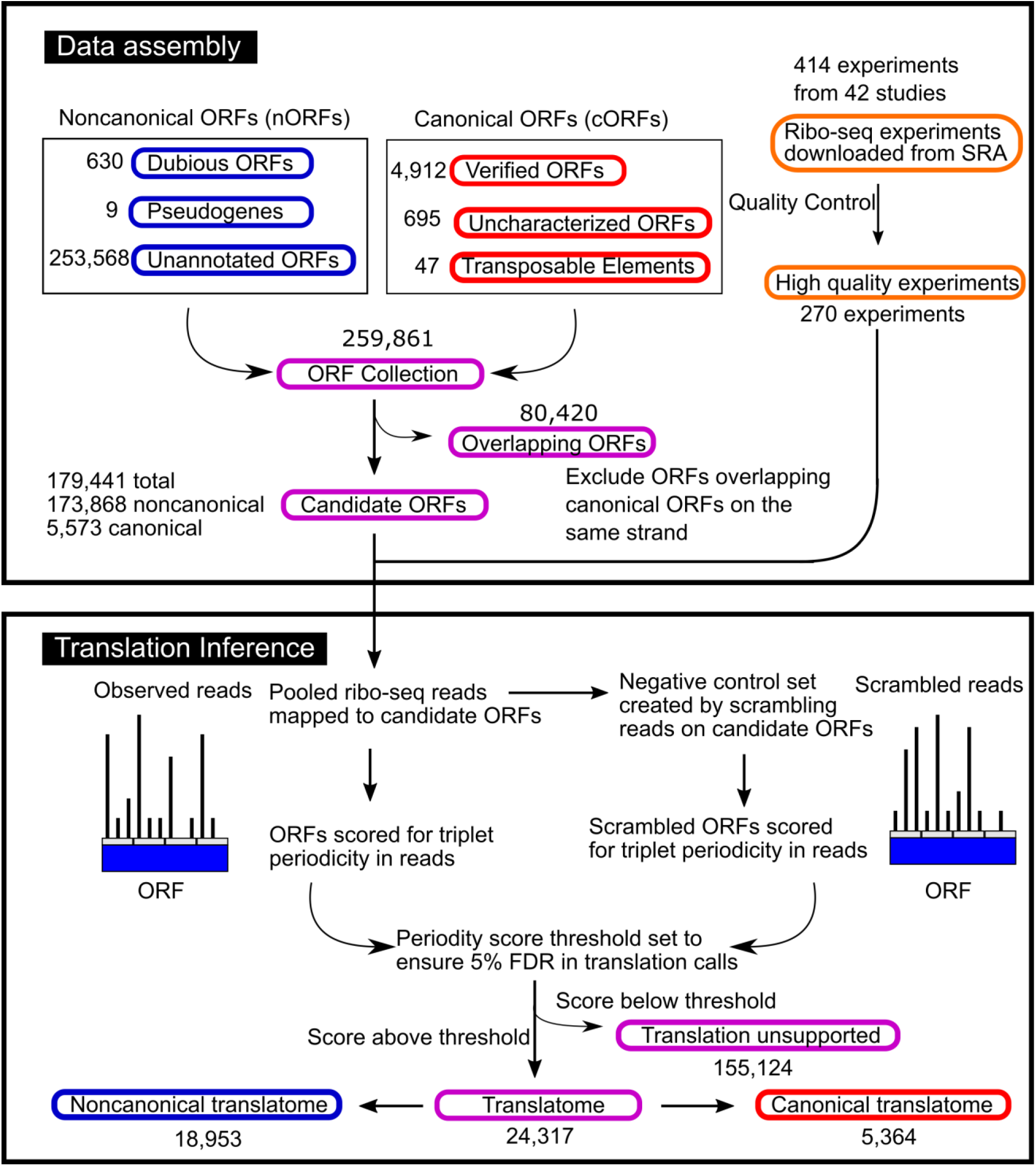
Workflow to identify translated ORFs in the *S. cerevisiae* genome using published datasets.

**Supplementary Figure 2:**
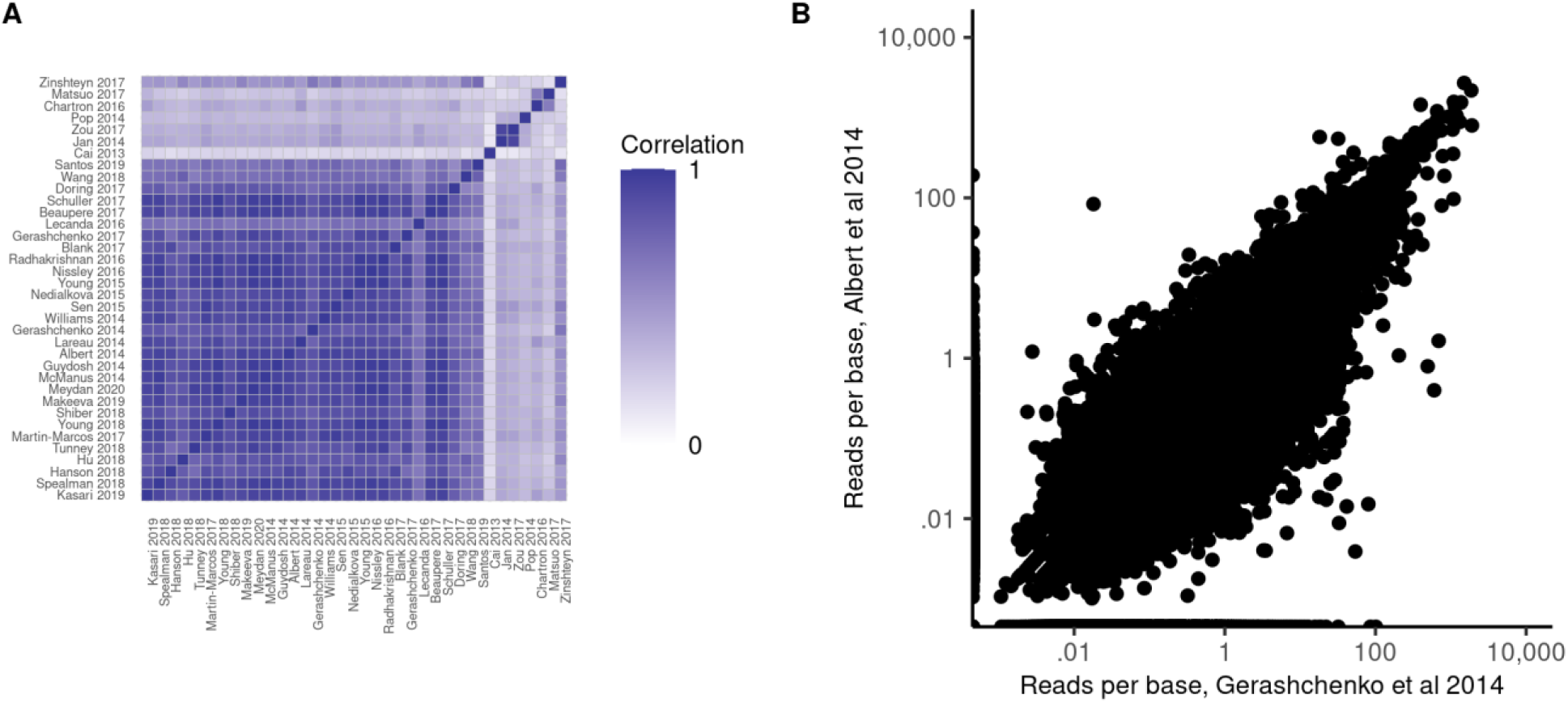
Translation patterns in candidate ORFs show high replicability between studies. A) Pairwise correlation between ribo-seq coverage of all candidate ORFs between studies included in the dataset. B) For each candidate ORF, the reads per base (considering only in-frame reads) are plotted for the two largest studies in the dataset.

**Supplementary Figure 3:**
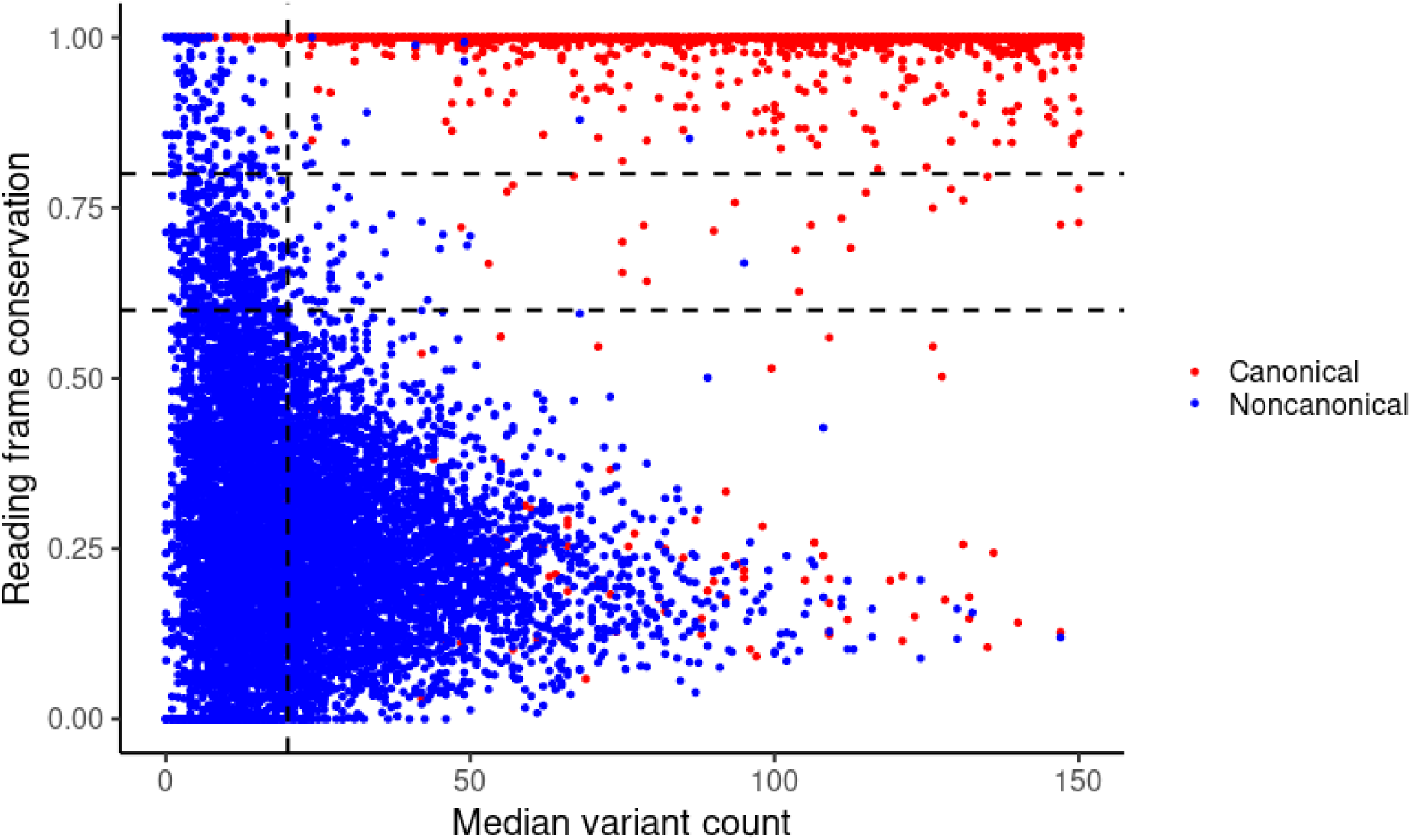
Nucleotide variation determines ability to distinguish conserved ORFs. Reading frame conservation for each nonoverlapping ORF is plotted against the median count of differences between the *S. cerevisiae* ORF and the aligned homologous sequence in each *Saccharomyces* relative. Colors indicate SGD annotation categories. To the right of the vertical line, there are two distinct populations separable by reading frame conservation; the intermediate region contains few ORFs. For ORFs to the left of the vertical line, with few differences in the ORF between species, there is no clear separation between high-RFC and low-RFC ORFs.

**Supplementary Figure 4:**
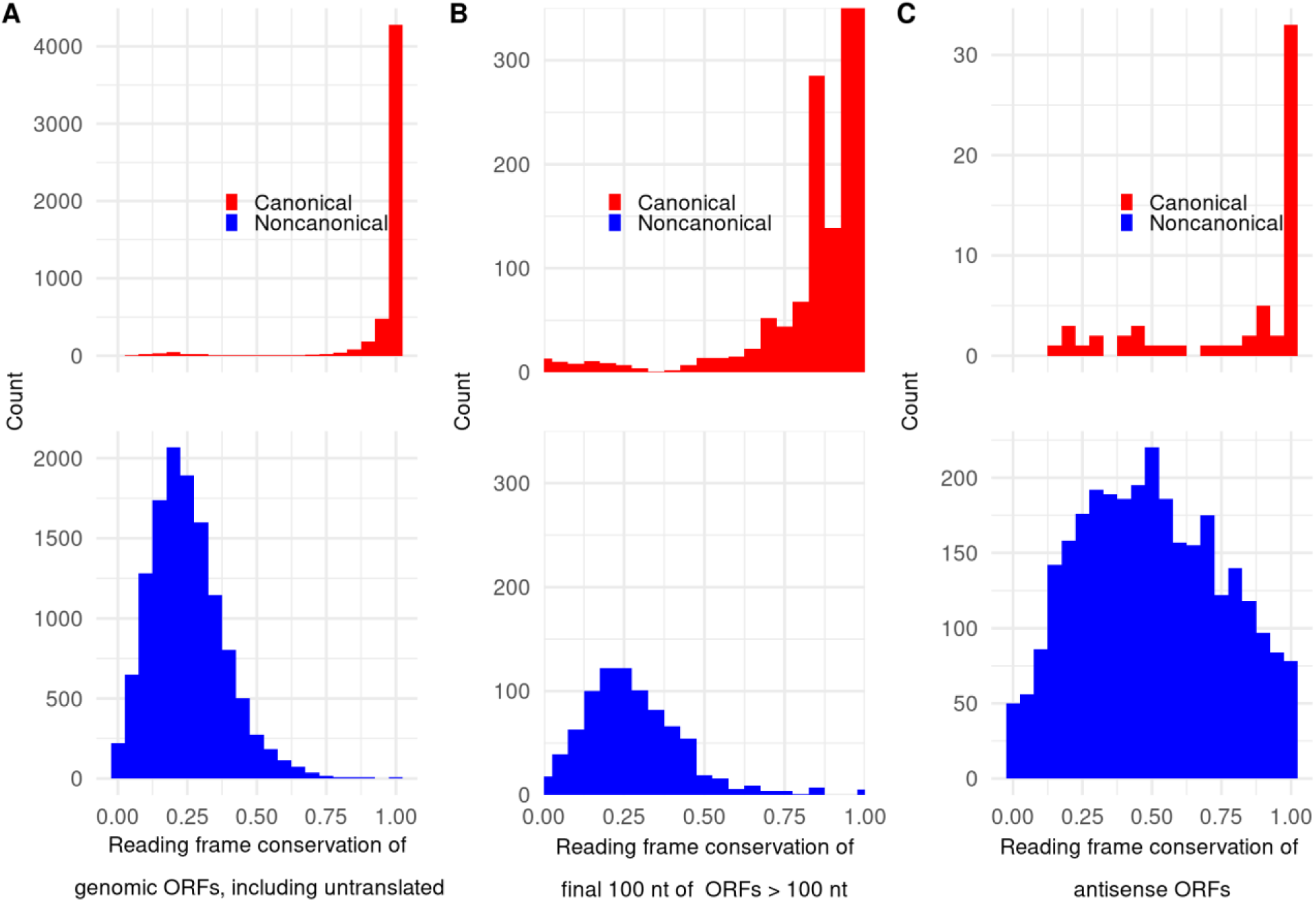
Distribution of frame conservation among classes of ORFs. A) The distribution of frame conservation among candidate ORFs in the genome, including both translated and untranslated ORFs. B) For all ORFs in the high information set at least 100 nt in length, RFC was calculated considering only the final 100 nt of the ORF. RFC was then plotted for both cORFs and nORFs. This was done to test whether low RFC in nORFs could be caused by inferring start codons upstream of the actual start codons for conserved nORF, which would lead to false inference of a low RFC value. However, the pattern considering only the final 100 nt is similar to the pattern observed for the full ORFs in Figure 4B, with a clear bimodal distribution, indicating that false start codon inference is likely not driving the pattern. C) The distribution of frame conservation is plotted for translated cORFs and nORFs that are antisense to canonical genes. In contrast to frame conservation among nonoverlapping ORFs, the distribution does not appear bimodal.

**Supplementary Figure 5:**
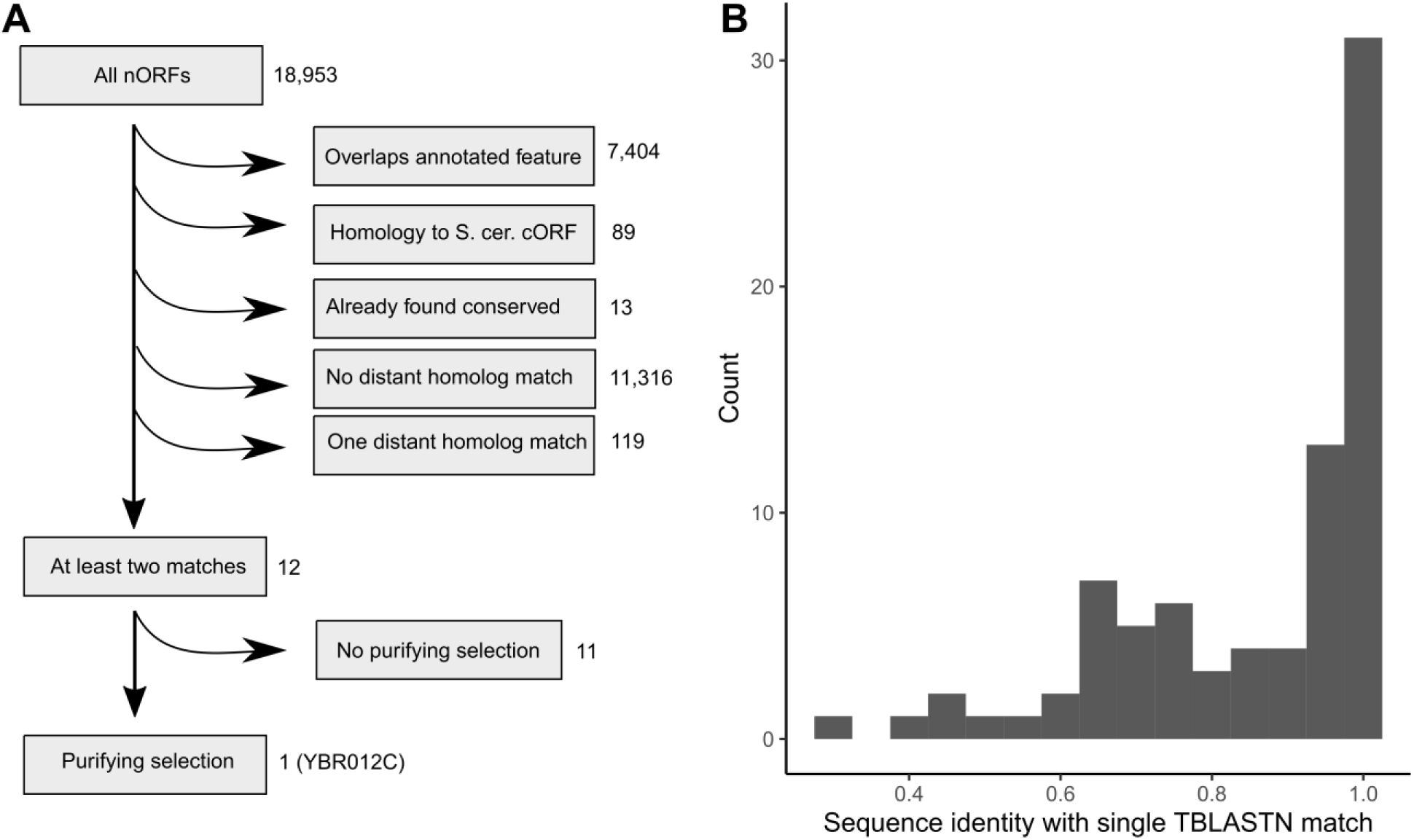
Identification of conserved genes in the noncanonical translatome using TBLASTN. A) Process for identification of conserved nORFs evolving under purifying selection. To be identified as conserved, an nORF could not overlap any annotated feature on the *S. cerevisiae* genome or have any homology to an *S. cerevisiae* cORF at a 10^-4^ BLASTP e-value threshold (as this makes BLAST results ambiguous) and have at least two identified homologs in a TBLASTN search at a 10^-4^ e-value threshold. Then, an additional indicator of selection was required (RFC > .8, or p-value < .05 in a test of neutrality using dN/dS or pN/pS). B) Among translated *S. cerevisiae* ORFs with a single TBLASTN hit (at a 10^-4^ e-value threshold) among budding yeasts outside the *Saccharomyces* genus, the distribution of sequence identities with that match is plotted. The existence of only a single match together with the prevalence of high sequence identities (>80%) suggests that the matches may be the result of genomic contamination rather than genuine homology, so at least two matches are required to accept homology as valid.

**Supplementary Figure 6:**
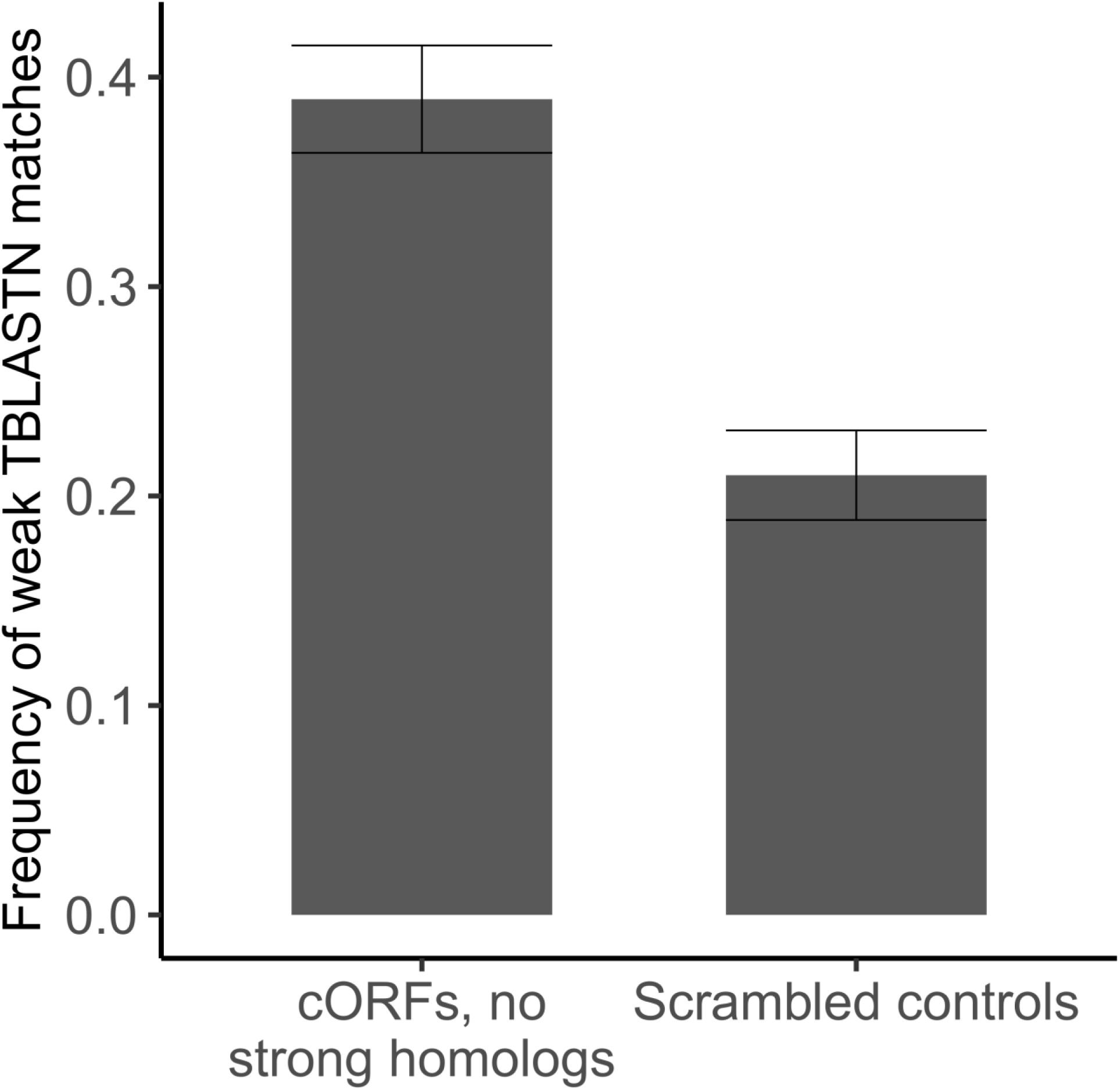
cORFs lacking high-confidence homologs are enriched in weak TBLASTN matches. The frequency of weak TBLASTN matches (10^-4^ < e-value < .05) among budding yeast genomes for cORFs that lack any strong matches, and controls consisting of the same sequences randomly scrambled. Error bars indicate standard errors estimated from bootstrapping.

**Supplementary Figure 7:**
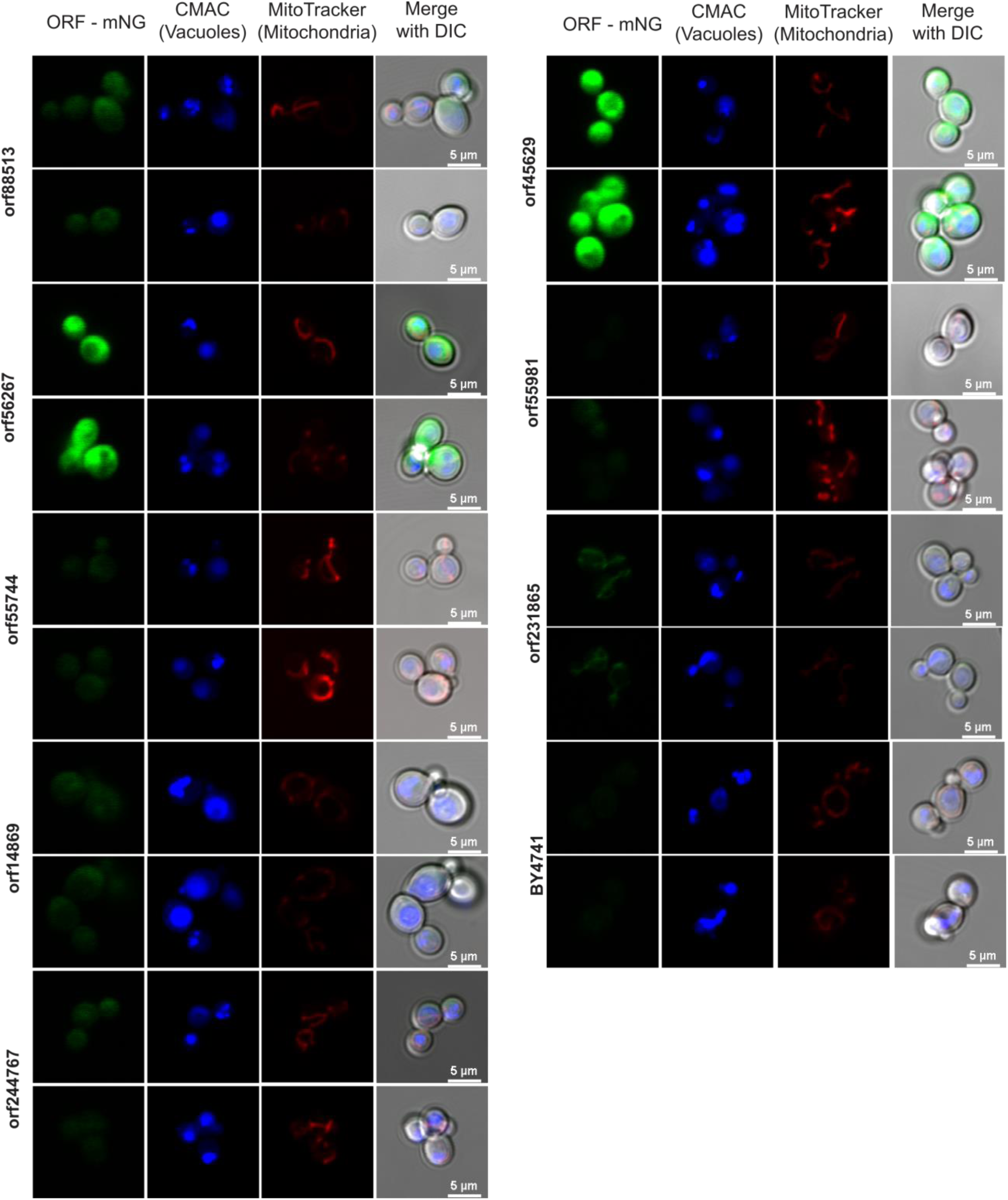
Microscopy of detected transient nORFs. Microscopy images of unannotated transient nORFs taken at 40X. Left panel show the expression of the nORFs tagged with mNeonGreen, middle panels the dyes CMAC Blue and MitoTracker Red for mitochondria and vacuoles identification, respectively, and the right panel the merge all the above channels with DIC. Two representative images are shown per strain; expression of orf55981 was not uniformly detected, with some cells showing expression and some not.

## Supplementary tables

**Supplementary Table 1:** Ribosome profiling experiments used for translation inference.

**Supplementary Table 2:** Ribosome profiling studies used for translation inference.

**Supplementary Table 3:** The yeast translatome.

**Supplementary Table 4:** Selection analysis of ORF groups in S. cerevisiae strains.

**Supplementary Table 5:** Phenotypes of canonical evolutionarily transient ORFs reported in literature.

**Supplementary Table 6:** Results of deletion mutant screen on transient nORFs using a gene replacement strategy.

**Supplementary Table 7:** Gene ontology analysis of genetic interactors of annotated transient ORFs.

**Supplementary Table 8:** Information on ORFs tested in translation inhibition experiment.

**Supplementary Table 9:** Strains used in this study.

## STAR Methods

### Key resources table

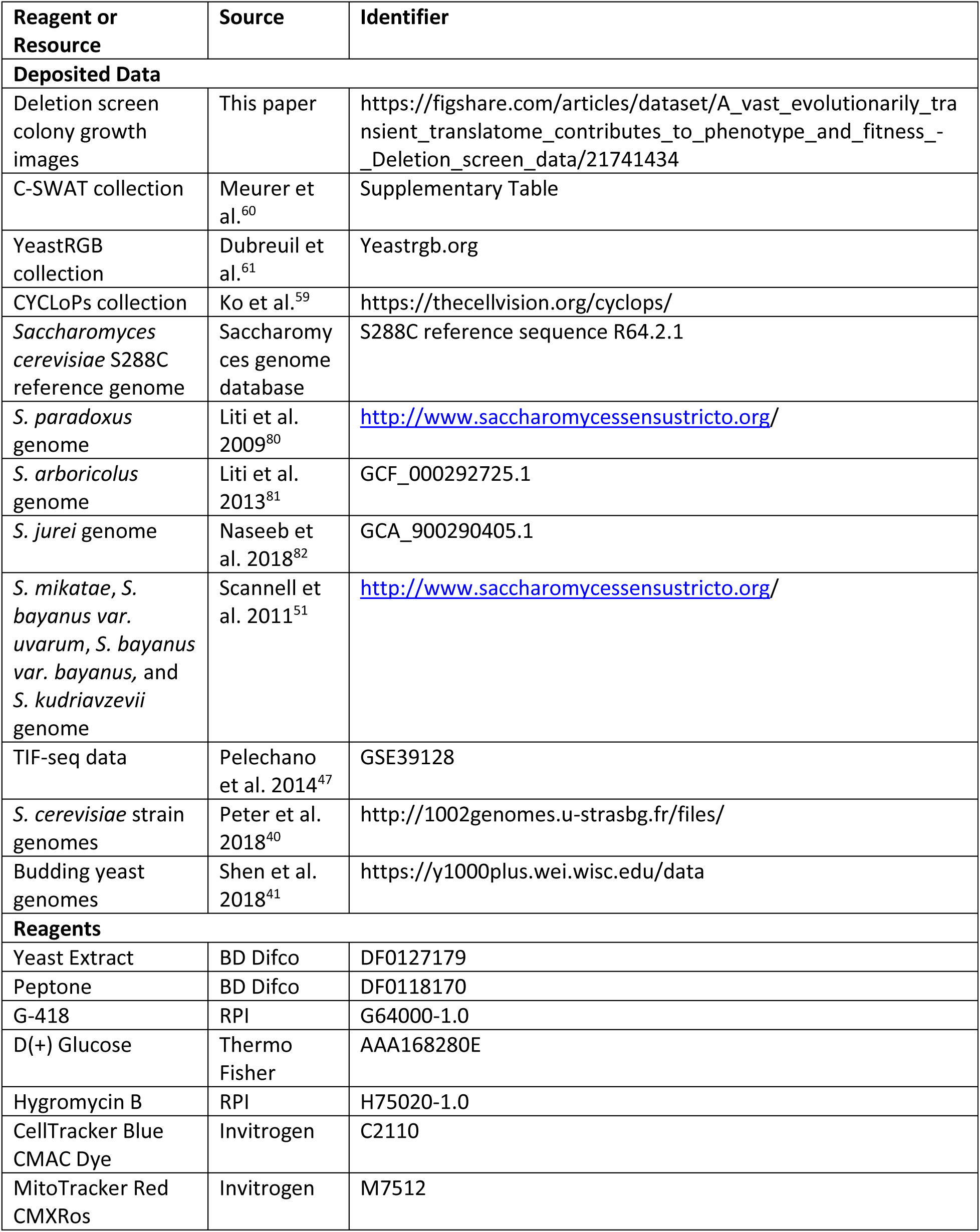
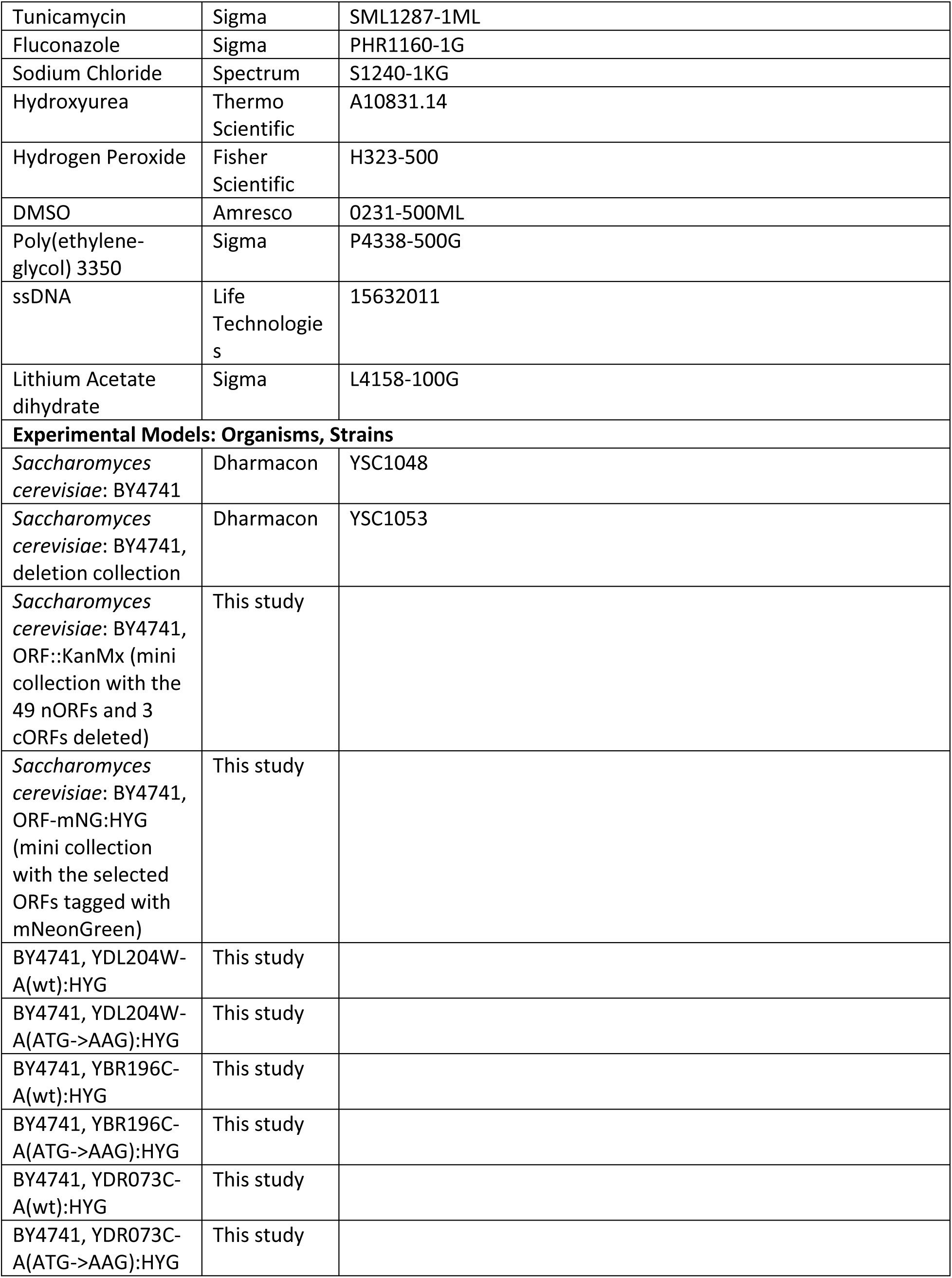
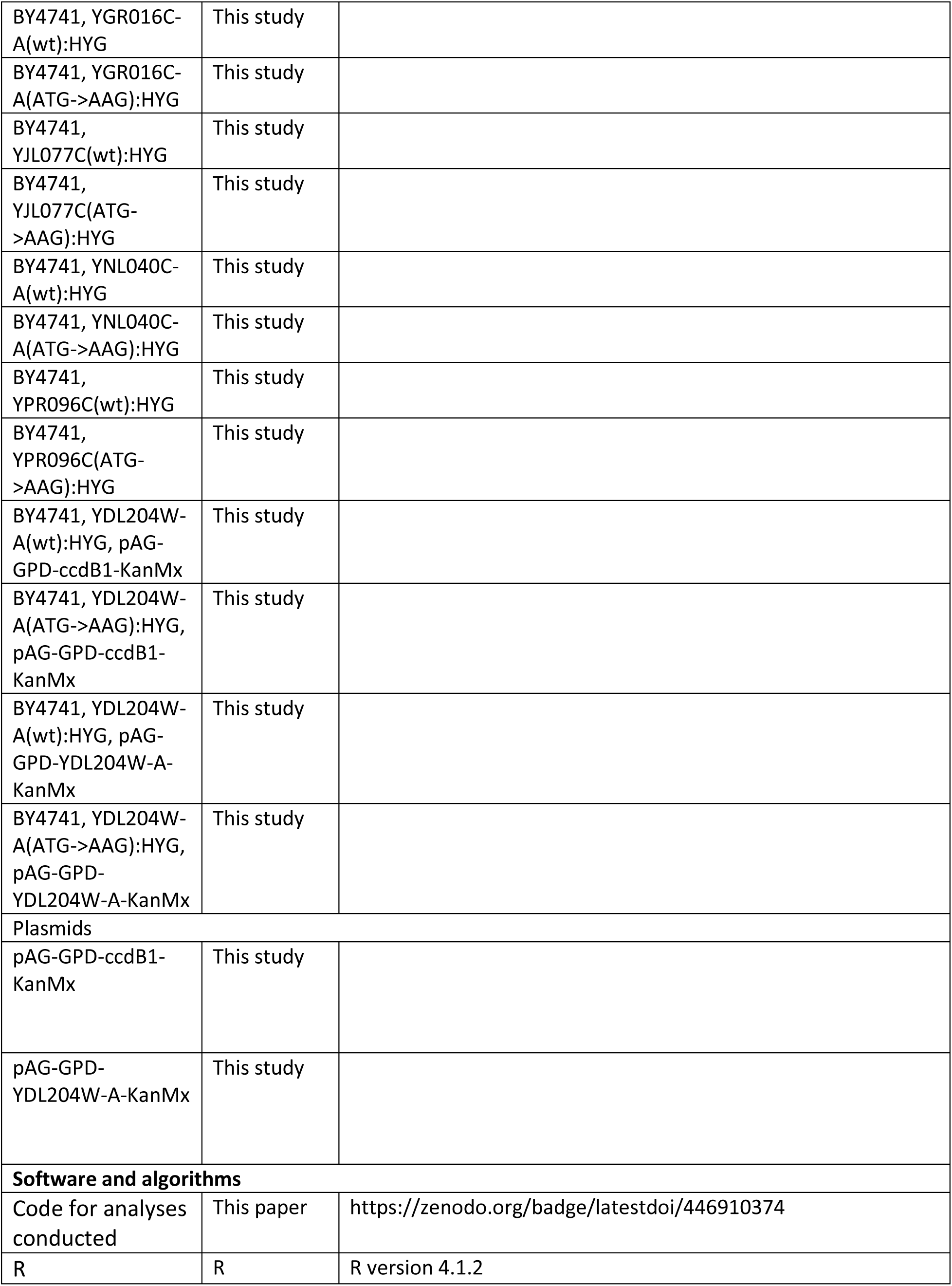
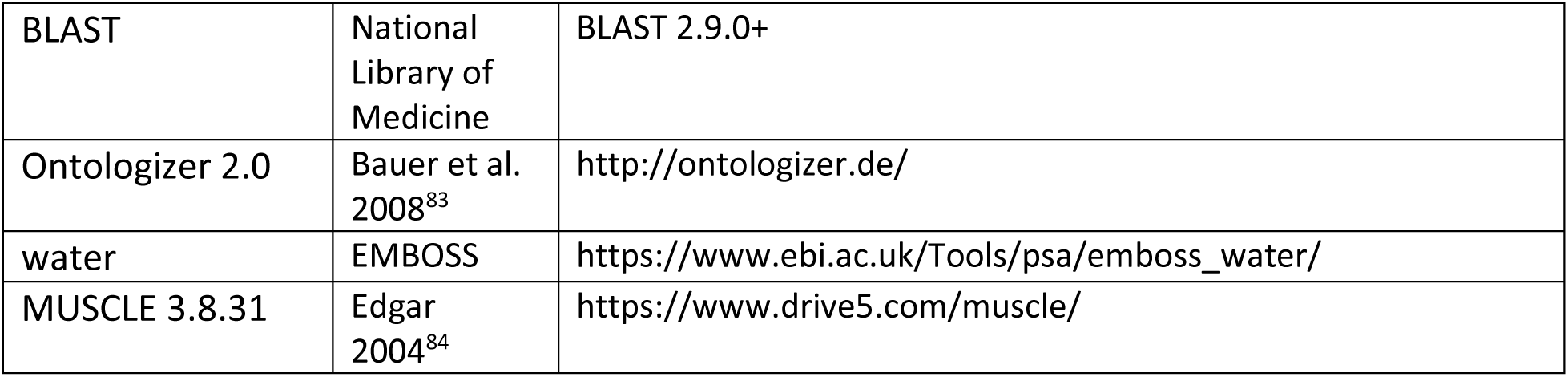

### Resources availability

#### Lead contact

Further information and requests for resources and reagents should be directed to and will be fulfilled by the lead contact, Anne-Ruxandra Carvunis.

#### Materials availability

All materials will be made available on request.

#### Data and code availability

• All original code has been deposited on GitHub and is publicly available as of the date of publication. DOIs are listed in the key resources table.
• Plate images for colony growth assays are available at Figshare and are publicly available as of the data of publication. DOIs are listed in the key resources table.
• Any additional information required to reanalyze the data reported in this paper is available from the lead contact upon request.

### Experimental Model and Subject Details

#### Yeast strains

All strains used in this study are derived from BY4741 (Dharmacon, YSC1048). The parental strain and all derivatives produced in this study are listed in Supplementary Table 9. The lithium acetate method^85^ was used to create new strains and selection was performed on appropriate selection plates. For genomic integration, the inserts were PCR amplified from plasmids or GBlocks.

### Method Details

#### Defining candidate ORFs

To identify a set of translated ORFs, a set of candidate ORFs was constructed for which translation status could be inferred using ribo-seq data. ORFs were identified on the R64.2.1 *Saccharomyces cerevisiae* genome assembly downloaded from SGD.^43^ The initial set of candidates consisted of all possible single-exon reading frames starting with an ATG, ending with a canonical stop codon, and having at least one additional codon between the start and stop. Among all ORFs that shared a stop codon, all but the longest were discarded. An ORF was considered canonical if it shared a stop codon with an ORF annotated as “verified”, “uncharacterized”, or “transposable element gene” on SGD. All other ORFs that overlapped a canonical ORF on the same strand were removed (including pairs of overlapping canonical genes) while ORFs that overlapped cORFs on the opposite strand were classified as antisense ORFs.

#### Yeast ribo-seq dataset collection and read mapping

A list of *S. cerevisiae* ribosome profiling (ribo-seq) studies was identified by conducting a broad literature search. For each study, all ribo-seq experiments were added to the dataset except those conducted on mutants designed to alter wildtype translation patterns. The full list of experiments and studies included is given in **Supplementary Tables 1** and **2**, respectively. The fastq files associated with each experiment were downloaded from Sequence Read Archive^86^ or the European Nucleotide Archive.^87^ If adaptors were present in the fastq file, they were trimmed. Reads were filtered to exclude reads in which any base had a Phred score below 20. For each remaining read, the number of perfect matches in the *S. cerevisiae* genome were identified, and only unique perfect matches were kept.

In initial mapping, reads were assigned to the genomic position aligning with the first base of the read. It was necessary to remap the reads such that the position assigned to the read instead corresponded to the first amino acid in the P-site of the translating ribosome, as in previous ribo-seq analyses^37^, so that the triplet periodic signal indicative of active translation overlaps precisely the bounds of translated ORFs. This was done by shifting all reads by the same number of positions, with the number determined separately for each read length and each experiment. To determine this number, a metagene profile was constructed: the number of reads in each of the -20 to +20 positions relative to the start codon was counted, accumulated over all annotated genes on Saccharomyces Genome Database (SGD)^43^. As there should be many more reads on the start codon of annotated genes than the sequence immediately upstream of these genes, the first attempt was to remap the first position with read count above a threshold to the first amino acid on the start codons, which then requires all other reads to shift by the same amount. The threshold selected was 5% of the total reads within 20 bases of the annotated start codons. The attempted shift was accepted if the expected triplet periodic pattern was obtained; i.e., there were more remapped reads on the first base of the codons of annotated genes than on the second or third base. Otherwise, a second shift was attempted from the next position exceeding the read count threshold, and so on until both criteria were met.

For quality control, presence of triplet periodicity was then tested for each read length in each experiment. The number of reads mapping (after remapping) to the first, second, and third position of each codon was counted among annotated genes, requiring at least twice as many reads in the first position than each of the second and third. If a read length failed this test for a given experiment it was excluded from further analysis, and if all read lengths for an experiment failed the experiment itself was excluded. All read lengths from 25 to 35 nucleotides were tested.

#### Translation calling

The iRibo program can be applied to any set of ribo-seq experiments to identify a set of ORFs with evidence of translation among those experiments. To construct a reference translatome, translation was inferred using ribo-seq data from the full set of experiments we collected that passed quality control (**Supplementary Table 3**). Separately, iRibo was also run on specific subsets of the full collection, including: experiments with or without the drug cycloheximide, experiments only on cells grown in YPD; only on cells grown on SD; and only on cells grown in YPD without cycloheximide (**Supplementary Table 3**). iRibo was also run separately for each individual study, generating lists of translated ORFs within each study.

Translation was assessed as follows: for each codon in each candidate ORF, the position within the codon with the most reads was noted, if any. The number of times each codon position had the highest read count across the ORF was then counted. The binomial test was then used to calculate a p-value for the null hypothesis that all positions were equally likely, against the alternative that the first position was favored. This p-value is an indicator of the strength of evidence for triplet periodicity favoring the first codon position.

To estimate the false discovery rate (FDR), a set of ORFs corresponding to the null hypothesis was constructed. For each ORF, the ribo-seq reads were scrambled randomly position by position (not read by read); e.g., if 10 reads mapped to the first base on the actual ORF, a random position in the scrambled ORF was assigned 10 reads, and so on. In this way the read distribution across positions was maintained but the spatial structure was eliminated. The same binomial test as used for the actual reads was then used on all scrambled-read ORFs. For every p-value threshold, the FDR can then be calculated as the number of scrambled ORFs with p-value below the threshold divided by the number of actual ORFs with p-values below the threshold. For each list of translated ORFs, the p-value threshold was set to give a 5% FDR among noncanonical ORFs; all ORFs with p-values below this threshold were then included in the translated set, whether canonical or noncanonical.

#### Estimating translation rates across different genomic contexts

All nORFs were partitioned into genomic contexts, with nonoverlapping nORFs classified by the relation between the nORF and any cORF located on the same transcript and antisense nORFs classified by partial or complete overlap of the opposite strand gene. The transcripts reported in Pelechano et al. 2014^47^ based on TIF-seq data were used for this analysis. An nORF was considered antisense if it overlapped an ORF annotated as “verified”, “uncharacterized”, “transposable element” or “blocked” on SGD on the opposite strand and nonoverlapping otherwise (ORFs overlapping annotated genes on the same strand were excluded from analysis, as described above). A nonoverlapping nORF was considered to share a transcript with a cORF or annotated non-coding RNA if any transcript fully contained both the nORF and the cORF or annotated RNA sequence; the ORF was then further classified as being either a uORF or dORF based on whether it was upstream or downstream of the cORF or RNA. If an nORF shared a transcript with both its upstream and downstream neighboring cORFs, it was classified according to the cORF that was closer.

#### Identifying homologous sequences of the *S. cerevisiae* ORF in other Saccharomyces genus species

Genomes were obtained from seven relatives of *S. cerevisiae* within the *Saccharomyces* genus: *S. paradoxus* from Liti et al. 2009^80^, *S. arboricolus* from Liti et al. 2013^81^, *S. jurei* from Naseeb et al. 2018^82^, and *S. mikatae*, *S. bayanus var. uvarum*, *S. bayanus var. bayanus,* and *S. kudriavzevii* from Scannell et al. 2011.^51^ Alignments were constructed between each *S. cerevisiae* ORF and its homologs in each *Saccharomyces* relative using synteny information. To identify anchor genes for syntenic blocks, BLASTP was run for each annotated ORF in *S. cerevisiae* against each ORF in the comparison species. Identified homolog pairs with e-value < 10^-7^ were selected as potential anchors. For each ORF in the *S. cerevisiae* genome, the upstream anchor *G0* and downstream anchor *G1* were selected that minimized the sum of the distance between the anchors in *S. cerevisiae* and the distance between the anchors in the comparison species; this sum was required to be less than 60 kb. The sequence between and including G0 and G1 were then extracted from both the *S. cerevisiae* genome and the comparison species and a pairwise alignment of the syntenic region was generated using MUSCLE 3.8.31.^84^

To confirm that the ORF was matched to genuinely homologous DNA, the alignment of the *S. cerevisiae* ORF along with its 50 bp flanking regions was extracted from the full syntenic alignment. The extracted region was then realigned using the Smith-Waterman algorithm^88^ with a match bonus of 5, a mismatch penalty of 4, and a gap penalty of 4. To test homology, 1000 alignments were constructed using the same score system in which the sequence of the comparison species was shuffled at random, reflecting a null hypothesis that the region was not homologous. The proportion of times the alignment of the real sequence scored better than the shuffled ones is a p-value indicating the strength of the null hypothesis against the alternative that the region is homologous. Homology was accepted as confirmed if the p-value was less than 1%, and alignments were excluded from analysis if homology was not confirmed.

If a syntenic alignment could not be constructed for a particular *S. cerevisiae* ORF and comparison species (because homology failed or there were no appropriate anchors), BLAST was attempted as an alternative method of finding the homologous DNA sequence. For these ORF sequences, BLASTn was run against the genome of the comparison species. For each reciprocal best matching pair with e-value < 10^-4^, the matched sequences in both species were extracted, together with a 1000 bp flanking region in both ends, and aligned using MUSCLE.^84^ DNA homology was then tested using Smith-Waterman alignment as described above.

#### Division of ORFs into high information and low information sets

Evolutionary analysis of ORFs was done separately for those ORFs for which there existed substantial information to test selection (“high information ORFs”) and those for which less information was available (“low information ORFs”). To be placed in the high information set, the ORF had to meet a homology criterion and a diversity criterion. The homology criterion required that DNA homology was confirmed in either a synteny or BLAST-based pairwise alignment with at least four other species in the *Saccharomyces* genus. For the diversity criterion, the number of single nucleotide differences (excluding gaps) was counted between the *S. cerevisiae* ORF and all its aligned sequence with confirmed homology among *Saccharomyces* genomes. The diversity criterion was satisfied if the median count of differences exceeded 20.

#### Reading frame conservation

Reading frame conservation is a measure of conservation of codon structure developed by Kellis et al. 2003^20^ and used here with some modifications. Calculation of reading frame conservation was done on a pairwise alignment of a genomic region containing the *S. cerevisiae* ORF (either a syntenic block between conserved genes or the 1000 bp flanking region around a BLAST hit). All single-exon ORFs (ATG to stop codon) in the comparison species were identified across this region. For each ORF in the comparison species, the reading frame conservation was calculated by summing up all points in the alignment where the pair of aligned bases are in the same position within the codon (i.e., both are in either the first, second, or third position) and dividing by the length of the *S. cerevisiae* ORF in nucleotides (including start and stop codons). Positions that align to gaps or are outside the range of the *S. cerevisiae* ORF are always considered to be not in the same codon position and do not add to the numerator. The ORF in the comparison species with the highest reading frame conservation is considered the best match, and the reading frame conservation of the *S. cerevisiae* ORF in relation to each other *Saccharomyces* species is defined as its reading frame conservation with its best match. In addition to the pairwise reading frame conservation of each *S. cerevisiae* ORF in relation to its homologs in all other species, an index of reading frame conservation (RFC) was defined equal to the average reading frame conservation of the *S. cerevisiae* ORF against all species in the *Saccharomyces* genus for which homologous DNA could be identified.

#### Detecting distant homology among *S. cerevisiae* ORFs

The genomes of 332 budding yeasts were taken from Shen et al. 2018.^41^ We applied TBLASTN and BLASTP for each *S. cerevisiae* translated ORF against each genome in this dataset (excluding the *Saccharomyces* genus). Default settings were used except for setting an e-value threshold of 0.1; results were then filtered by a stricter e-value threshold as described in each analysis. The BLASTP analysis was run against the list of protein coding genes used in Shen et al. 2018^41^ while the TBLASTN analysis was run against each entire genome. In the TBLASTN analysis, scrambled sequences of each *S. cerevisiae* ORF were also included as queries to serve as a negative control.

#### Tests of selection using the dN/dS and pN/pS ratios

Variant call file data for 1011 *S. cerevisiae* isolates was taken from Peter et al. 2018.^40^ For each ORF, nucleotide diversity was estimated from the full set of isolates. Nucleotide diversity was estimated as the mean number of differences per site in the ORF between any pair of isolates. To calculate dN/dS, the consensus sequence among all isolates was determined. At each position in the consensus, the three possible nucleotide variations were recorded as possible polymorphisms and distinguished by polymorphism type (12 possible combinations of consensus and variant nucleotide) and whether they would result in a synonymous or nonsynonymous difference from the consensus. If at least one isolate had the polymorphism, the polymorphism was also recorded as observed. All possible and observed polymorphisms were counted among all considered ORFs.

The pN/pS ratio was calculated in a similar manner to Ruiz-Orera et al. 2018^28^ and could be applied to either a single ORF or a group of ORFs. For each ORF under consideration, the consensus sequence among all isolates was determined. At each position in the consensus, the three possible nucleotide variations were recorded as possible polymorphisms and distinguished by polymorphism type (12 possible combinations of consensus and variant nucleotide) and whether they would result in a synonymous or nonsynonymous difference from the consensus. If at least one isolate had the polymorphism, the polymorphism was also recorded as observed. All possible and observed polymorphisms were counted among all considered ORFs.

Consider a variant *X*→ *Y* where *X* is the consensus at a site and *Y* is a possible variant. The probability of observing variant *Y* at a position with consensus *X*, *px*→*Y* was estimated as the observed count of X→Y variant sites divided by the possible count of X→Y variant sites. Under neutrality, the expected count of either synonymous or nonsynonymous X→Y variant sites is then the product of *px*→*Y* and the number of possible synonymous or nonsynonymous X→Y variant sites. In this manner the expected and observed counts of synonymous and nonsynonymous variants were calculated. The pN/pS ratio is then estimated as:

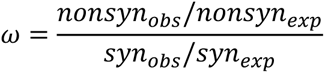

Under neutrality, then, the expected count of X→Y nonsynonymous variant sites is the number of possible such variant sites times the expected probability of this variant. In this manner the expected and observed counts of all synonymous variant types were calculated. To test for deviation from neutrality, we used a chi-squared test with one degree of freedom to compare observed vs. expected counts of synonymous and nonsynonymous variants. Standard errors for the pN/pS ratio in group analyses were estimated by bootstrapping: the ORFs in the group were resampled with replacement 1000 times and the pN/pS ratio was calculated each time. The standard error was then estimated as the sample standard deviation among the 1000 pN/pS ratios.

The dN/dS ratio was calculated based on differences in the pairwise ORF alignments *S. cerevisiae* and its closest relative *S. paradoxus*. Each S. cerevisiae ORF was associated with an *S. paradoxus* ORF for which the pair had the highest reading frame conservation (or none if homology with *S. paradoxus* was not confirmed or the highest reading frame conservation was 0). Counts of differences were made only for codons that shared the same frame between these ORFs and with at most one nucleotide difference between the codons. For every eligible position in the *S. cerevisiae* ORF, each possible *S. paradoxus* difference was counted and distinguished by whether the difference was synonymous or nonsynonymous and by type (four *S. cerevisiae* nucleotides, each with three possible *S. paradoxus* differences). These observed and possible differences were then used to estimate the dN/dS ratio in the same way as described above for the pN/pS ratio.

Among nORFs with high RFC, the strong conservation in *Saccharomyces* permitted calculation of dN/dS over the entire *Saccharomyces* tree, and so this was done as an additional test of selection (as reported in Table 1). For this analysis, ancestral reconstruction of the *Saccharomyces* phylogeny was conducted using PRANK^89^ with parameters -showanc -showevents -once -prunetree -keep. Ancestral reconstruction included all species in which DNA homology was confirmed. Codons were only used for counting substitutions if they shared frame conservation among all species. Observed and possible substitutions were counted across each branch and distinguished by substitution type and whether the substitutions were synonymous or nonsynonymous. Then, dN/dS was estimated in the same way as described for pN/pS above.

### Classification of ORFs into transient and conserved sets

All high-information nonoverlapping translated ORFs with RFC > 0.8 were classified as conserved (**Figure 4A**). An nORF was also classified as conserved if it overlapped no annotated feature on SGD, had TBLASTN matches with e-value < 10^-4^ with at least two species outside the *Saccharomyces* genus and showed at least one additional signature of purifying selection (RFC > 0.8 or a p-value < 0.05 in a test of neutrality using dN/dS or pN/pS) (**Supplementary Figure 5A**).

Nonoverlapping ORFs were excluded from classification in the transient set if they showed homology to an ORF classified as conserved in *S. cerevisiae* (e-value < 10^-4^ using BLASTP) or to any sequence among budding yeasts outside *Saccharomyces*^41^ (e-value < 10^-4^ using TBLASTN). Among remaining translated ORFs, all high-information ORFs with RFC < 0.6 were classified as transient. Low information ORFs were divided into groups and classified as transient if no group they belonged to showed evidence of selection in dN/dS analysis, pN/pS analysis, or weak homology matching analysis. Two low-information groups were cORFs and antisense nORFs. Low information nonoverlapping nORFs were each assigned to three groups corresponding to deciles of translation rate, coding score and ORF length. Analyses of dN/dS and pN/pS are described above. For weak homology detection, the number of ORFs with at least two weak TBLASTN matches (e-value < 0.05) to budding yeast genomes collected by Shen et al. 2018^41^ (excluding *Saccharomyces* species) was counted for both actual and scrambled ORF sequences. Selection was inferred if actual matches significantly (p < 0.05) exceeded scrambled matches using Fisher’s exact test. Only ORFs that did not overlap any annotated feature on SGD were included in weak homology detection analysis.

#### Coding score calculation

The coding score, described by Ruiz-Orera et al. 2014^90^, is a measure of how close the hexamer (i.e., the nucleotide sequence of a pair of adjacent codons) frequency of an ORF is to the hexamer of coding vs. noncoding sequences. Higher scores indicate a more gene-like hexamer distribution. Coding hexamer frequencies were calculated among all ORFs annotated as “verified” or “uncharacterized” by Saccharomyces Genome Database.^43^ Noncoding hexamer frequencies were calculated for all intergenic sequences (sequences in between verified or uncharacterized ORFs) in the *S. cerevisiae* genome. As intergenic sequence has no codon structure, hexamer frequencies for intergenic sequence were counted as if read in each possible coding frame. The score was then calculated as described in Ruiz-Orera et al. 2014.^90^

#### Identification to transient ORFs with detectable translation products in published microscopy studies

Published results were examined from fluorescent tagging experiments where the expression of ORFs was driven by native promoters and terminators. A list of ORFs detected in 15 GFP-tagged screens on wildtype strains in either normal conditions or with chemical treatment (hydroxyurea or rapamycin) were retrieved from the CYCLoPs database.^58, 59^ Lists of ORFs detected in the C-SWAT tagging library were taken from Meurer et al. 2018^60^ and from YeastRGB^61^. ORFs with fluorescent intensity below the reported detection threshold in each screen were filtered out. Transient ORFs that showed detectable translation products in at least one screen were considered as detected.

#### Literature analysis of transient translatome cORFs

For each transient translatome cORF, we examined all publications listed on SGD as “primary” or “additional” literature for the ORF. If the ORF had a phenotype in any listed publication, we noted the evidence for the phenotype (**Supplementary Table 5**).

#### Genetic interaction analysis

Single mutant fitness and genetic interaction data were downloaded from TheCellMap.org.^91^ In this dataset, mutants of nonessential genes are full deletions and mutants of essential genes are temperature-sensitive alleles. Transient ORFs were all nonessential. Different temperature-sensitive alleles for the same essential gene were treated separately. We removed all genes or transient ORFs with a genomic overlap to another genetic element from our analyses as it is not possible to assign the observed phenotypes to either of the overlapping pairs.

We counted the number of transient ORF and nonessential genes that showed at least one genetic interaction with Ɛ<-.2 and p-value < 0.05 (a negative genetic interaction) or Ɛ<-.35 with a p-value<0.05 (a synthetic lethal interaction). We then divided this number by the total number of transient ORFs or nonessential genes in the Costanzo et al. 2016^69^ genetic interaction dataset to calculate the percentage showing at least one genetic interaction. We used Fisher’s exact test to assess the significance of differences between percentages of nonessential genes and transient ORFs.

Gene ontology analysis of the interactors of each ORF was conducted with Ontologizer^83^, using Benjamini-Hochberg multiple testing correction and the term-for-term calculation method. The gene association file was downloaded from SGD. Gene ontology evidence codes relating to genetic interactions (IGI and HGI) were not used.

#### Creation of yeast strains

Deletion mutant strains for 49 transient nORFs and 3 transient cORFs were created by using homologous recombination to replace the ORFs with a KanMX cassette. Transformations were done using the LiAc/PEG protocol^85^ in the background BY4741 strain, and selected in media containing G-418. After an initial screen of these strains, a subset of the deletion strains that showed strong deleterious effects were transformed a second time, also using the LiAc/PEG protocol^85^, to replace the KanMx cassette with either an intact copy of the original ORF, or a mutant copy of the ORF with the start codon ATG and (in some cases) additional in-frame ATG codons mutated to AAG to prevent translation. This was accomplished by using homologous recombination to replace the KanMx cassette with a construct containing the intact or mutant ORF followed by a hygromycin resistance cassette. These constructs were synthesized by IDT (Integrated DNA Technologies). The resulting transformants were selected in agar plates containing hygromycin. All positive clones were sequenced to confirm presence of either the restored wildtype ORF or the ORF with a mutated start codon.

Strains containing an mNeonGreen tag for microscopy purposes were also made by homologous recombination using the LiAc/PEG protocol^85^ in the BY4741 background. The mNeonGreen and hygromycin cassette sequences were amplified from a plasmid using primers containing homology to the 3’ of each ORF. The primers were designed to remove the STOP codon of each ORF and place the mNeonGreen in frame with the ORF, to be expressed under its native promoter. Positive clones were selected on agar plates containing hygromycin.

All strains were kept in glycerol stocks at −80 °C in 96 and 384-well format until used for screening. Strain genotypes are listed in **Supplementary Table 9**.

#### Screening strategy for fitness estimation

Both rounds of deletion screening were conducted at 1536 colony density, with 1 in 4 colonies on the plate being reference strains used to correct for spatial biases as described in Parikh et al. 2021.^92^ In the initial deletion screen, each mutant strain was tested using 12 replicates; 72 replicates were tested per strain in the start codon mutant screen. Conditions tested were YPDA and YPDA+DMSO as unstressed conditions and five stress conditions: YPDA supplemented with 1M NaCl, 100mM Hydroxyurea, 0.6µM Tunicamycin, 25µg/ml Fluconazole, or 30mM Hydrogen peroxide (H2O2). Agar plates were incubated and imaged periodically until the colonies reached saturation. The plate handler Singer ROTOR (Singer Instrument Co. Ltd) was used to prepare all plates starting from glycerol stocks. Serial imaging of the plates was conducted using the spImager Automated Imaging System (S & P Robotics Inc., Ontario, Canada). The images were analyzed in bulk using a custom script made using functions from the MATLAB Colony Analyzer Toolkit^92^ to provide colony size estimations (https://github.com/sauriiiin/lid_personal/blob/master/justanalyze.m). The output files containing colony size information along with the images is available at https://bit.ly/3xtzHJO. The LI Detector analytical pipeline^92^ was used to correct for spatial biases in colony size and obtain colony fitness estimates. Strain fitness was estimated as the median of bias-corrected colony size among replicates of the strain at 40 hours in the initial screen and 90 hours in the start codon mutant screen. In the LI Detector pipeline^92^, sets of reference colonies are treated as if they were replicates of a mutant strain, with their median fitness calculated in order to construct an empirical null distribution of median fitness values to compare with estimated strain fitness. Strains were called as beneficial or deleterious using a 5% false discovery rate threshold based on this empirical null distribution. For any selected fitness threshold used to infer deleterious strains, the false discovery rate can be calculated as the proportion of null distribution fitness values below that threshold divided by the proportion of mutant strain fitness values below the threshold. Thus, fitness thresholds were selected such that a 5% FDR was obtained and strains with fitness below that threshold were inferred to be deleterious. In the same manner, a list of beneficial strains at 5% FDR was also selected.

#### Liquid growth assay

For liquid growth assays, cells were first grown in liquid YPDA media overnight at 30°C in a 96-density microplate. These were then used to inoculate a new 96-density microplate with 150μl YPDA+ stress conditions (1M NaCl, 100mM Hydroxyurea)) using the Singer ROTOR (Singer Instrument Co. Ltd). This microplate was incubated at 30°C with constant double orbital shaking for a period of 72h on microplate reader Biotek Synergy H1 (Aligent Technology Inc.). Optical density readings at 600nm (OD_600_) were taken every 15 minutes.

#### Microscopy

The strains containing the ORFs tagged with mNeonGreen were imaged on a Nikon TiE2 inverted A1R confocal microscope. A first screening was done at high density in 96-well plates with a 40x water objective, to assess the success of the transformations. Plates were incubated with CellTracker Blue CMAC Dye (Invitrogen) and MitoTracker Red CMXRos Dye (Invitrogen) at least 10 min prior to imaging. Plates were then imaged in 4 channels (405, 488, 561, and DIC), and 3 fields of view were taken for each strain that contained many cells. Strains that demonstrated visibly higher signal in the green channel (488nm) compared to a non-transformed background strain were selected to examine in single dishes under a 100X oil objective to more accurately evaluate sub-cellular localization. All strains were imaged in triplicate at high density and triplicate in dishes (once without CMAC and MitoTracker and two times with the dyes).

### Quantification and statistical analysis

Statistical analyses were performed in R version 4.1.2. Details for each statistical test and analysis can be found in the results section and figure legends.

## Notes

### Summary of Updates

Many analyses have been updated and new experiments conducted and described.

